# Wasp controls oriented migration of endothelial cells to achieve functional vascular patterning

**DOI:** 10.1101/2020.09.15.296574

**Authors:** André Rosa, Wolfgang Giese, Katja Meier, Silvanus Alt, Alexandra Klaus-Bergmann, Lowell Edgar, Eireen Bartels, Russell Collins, Anna Szymborska-Mell, Baptiste Coxam, Miguel O. Bernabeu, Holger Gerhardt

**Author notes:** These authors contributed equally to this work (Wolfgang Giese, Katja Meier).

## Abstract

Endothelial cell migration and proliferation are essential for the establishment of a hierarchical organization of blood vessels and optimal distribution of blood. However, how these cellular processes are coordinated remains unknown. Here, using the zebrafish trunk vasculature we show that in future veins endothelial cells proliferate more than in future arteries and migrate preferentially towards neighboring arteries. In future arteries endothelial cells show a biphasic migration profile. During sprouting cells move away from the dorsal aorta, during remodelling cells stop or move towards the feeding aorta. The final morphology of blood vessels is thus established by local proliferation and oriented cell migration to and from neighboring vessels. Additionally, we identify WASp to be essential for this differential migration. Loss of WASp leads to irregular distribution of endothelial cells, substantially enlarged veins and persistent arteriovenous shunting. Mechanistically, we report that WASp drives the assembly of junctional associated actin filaments and is required for junctional expression of PECAM-1. Together, our data identify that functional vascular patterning in the zebrafish trunk utilizes differential cell movement regulated by junctional actin, and that interruption of differential migration may represent a pathomechanism in vascular malformations.

## Introduction

The formation of a functional network of blood vessels with precisely controlled hierarchy and morphology is a crucial process during embryogenesis, development and disease progression. Virtually all blood vessels derive from angiogenesis, a process that entails the tight coordination of several endothelial cell (EC) behaviors including migration, proliferation and lumen formation^1–3^. After the establishment of sprout connections^4^, blood vessel morphogenesis and remodelling is accomplish by endothelial oriented migration and proliferation within established vessels, in response to the local needs of tissues^5^ as well as hemodynamic forces ^6–9^.

Endothelial cell migration has been extensively studied in different *in vivo* and *in vitro* systems^10^ as well as the signals that potentiate cell proliferation and how both these processes can contribute to vascular growth^11,12^. Nevertheless, as most studies investigated tip and stalk cell processes in the initial sprouting process as well as during the establishment of the primitive plexus, a detailed understanding of quantitative endothelial dynamics in vascular remodelling is lacking. Therefore, to develop new models that predict the cellular behaviors that control vascular remodelling and hierarchical vascular patterning, a detailed quantitative characterization of EC dynamics and vascular morphology in this context will be required.

Although many aspects of cell migration and proliferation rely on the control of the actin cytoskeleton^13,14^ the exact role of actin regulators and in particular of nucleators and nucleation promoting factors (NPF) is not well understood in the context of vessel remodelling. Indeed several studies have shown that inhibition of F-actin polymerization compromises the process of elongation and rearrangement of cells^15,16^ and carefully controlled, inhibition of actin polymerization has been highly informative on the role of cellular projections like filopodia and cell movement^17^. Nevertheless, as many of these studies took advantage of powerful toxins that specifically destabilize actin polymerization or interfere with acto-myosin contractility, we lack deeper insight into the molecular mechanism required for F-actin regulation driving selective cellular processes in vascular remodelling.

Here, using the trunk vasculature of the zebrafish as a model, we performed a detailed quantitative analysis of EC dynamics throughout initial sprouting, lumenisation, anastomosis and remodelling of intersegmental vessels (ISVs) into a balanced network of arteries and veins. Bespoke computational analysis shows that immigration and emigration of cells between vessels rather than local proliferation determines the diameter of individual segments. The general direction of movement hereby is from veins towards arteries, with decreasing speed as vascular remodelling progresses. We identified the actin nucleation promoting factor WASp, previously thought to be selectively expressed in cells of hematopoietic origin, as a critical regulator of this behavior, in part by establishing junction associated actin structures thought to transmit forces that drive cell intercalation^18^. Loss of WASP function in vivo drives vascular malformations, suggesting that loss of directional collective migration responses may contribute to vascular pathologies.

## Results

### EC directed migration depends on vessel specification

ECs can migrate either individually or collectively as a group of cells in response to guidance cues^19^. The zebrafish trunk vasculature develops as the result of collective migration, where sprouts originating from the dorsal aorta, equally distanced by the length of one somite, migrate dorsally each led by a tip cell and followed by stalk cells^20^. To better understand the cellular and molecular processes that contribute to vessel morphogenesis in the zebrafish trunk, we took advantage of endothelial specific reporter lines that label the cell nucleus and filamentous actin (*fliep:nls-mCherry; fliep:lifeactGFP*, respectively)^17,21^. We captured time-lapse image sequences using a spinning disk confocal microscope to follow the development of blood vessels from 26 to 44 hours post-fertilization (hpf). At 26 hpf arterial sprouts (sprouts coming from the dorsal aorta) are composed of an average of three ECs migrating dorsally (Figure 1 A timestamp at 26 hpf). Around 28-30 hpf neighboring arterial sprouts fuse through anastomosis forming the dorsal longitudinal anastomotic vessel (DLAV) (Figure 1 A, timestamp at 28:30h PF). At 31 hpf, venous sprouts emerge from the posterior cardinal vein (PCV), connecting to existing arterial ISVs. The remodelling of these connections between vessels (Figure 1 A, timestamp at 31:00-35:20h PF) ultimately leads to the formation of a balanced network of arterial and venous ISVs (aISVs and vISVs) by 44 hpf (Supp Movie 1).

**Figure 1:**
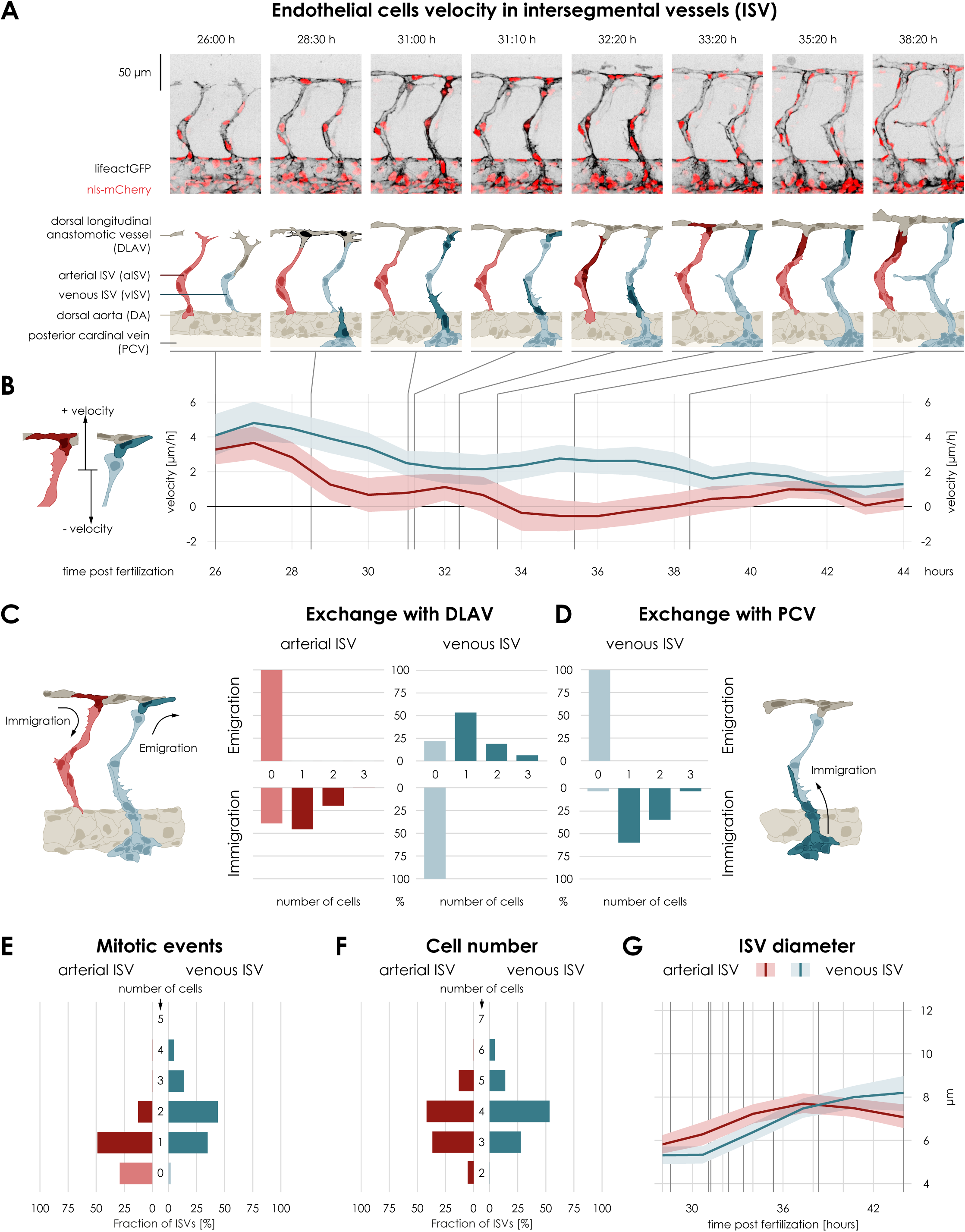
EC velocity changes with dynamic cellular rearrangements: **A**, Montage of two ISVs sprouts from 26 hpf until 38 hpf. Top panel: Live imaging of Tg[*fliep:nls-mCherry; fliep:lifeactGFP*] Black: F-actin, Red: nuclei. Bottom panel: Pictogram of live imaging; red ECs represent cells in aISV, blue ECs represent cells in vISV. Dark red ECs indicate cell division events (timepoint 32:20) and immigration of cells from DLAV to aISV (timepoints 33:20, 35:20-38:20). Dark blue cells indicate secondary sprouts (timepoint 28:30), vascular remodelling (timepoint 31:00, most ventral EC), cell division events (timepoint 31:00, most dorsal ECs) and emigration of ECs towards the DLAV (timepoint 31:10, 33:20-38:20) and immigration of ECs from PCV (timepoint 31:00-32:20). Dark grey cells represent cell anastomosis (timepoint 28:30). **B**, Collective EC velocity (μm/h) across time post fertilization (hours). Red: velocity for ECs in aISV, blue: velocity of ECs in vISV. Left pictogram demonstrates the referential for positive and negative velocity relative to dorsal and ventral movement. Data are mean ±CI (95%). **C**, Number of cells that exchange from ISVs with DLAV and fractions of ISVs that are affected. **D**, Number of cells that exchange from PCV with vISV and corresponding fractions of vISVs. **E**, Number of mitotic events in ISVs. **F**, Cell number in ISVs 44 hpf and distribution among ISVs. **G**, Diameter size (μm) across time post fertilization (hours) for aISV (red) and vISV (blue). Black lines represent the equivalent time points in montage. Data are mean ±CI (95%).

We systematically analyzed the movement of individual cells in a set of developing intersegmental vessels in a 10-somite region (Suppl. Figure 1 and see: Computational analysis pipeline in methods section) from a collection of 14-25 zebrafish embryos (Suppl. Figure 2). Orientation of cell migration was defined relative to the ventral-dorsal movement of cells in sprouts and nuclei movement was used as a proxy for cell migration. Nuclei moving dorsally were considered to migrate with a positive velocity, whereas nuclei with ventral displacement were assigned negative velocity (Figure 1B). The distinction between nuclei of sprouts, after DLAV formation, that will remodel into a vISV (future vISV) and nuclei from sprouts that will become an aISV (future aISV) was made *a posteriori*, i.e. we observed the remodeled network at 44 hpf and rewind the time-lapse sequence back to 26 hpf, marking each ISV as arterial or venous according to their final identity.

At 27 hpf the average cell velocity of ECs reached a maximum of 5 μm/h in sprouts of future veins, and 3.9 μm/h in sprouts of future arteries (Figure 1 B). Given that we assign the arterio-venous identity of the vessels at the endpoint of the time-lapse analysis, for the sake of simplicity, we will in the following refer to venous and arterial ISVs even when describing developmental stages at which this identity is not yet established. Between 28 and 31 hpf, arterial sprouts begin to anastomose with their ipsilateral neighbors to form the dorsal longitudinal anastomotic vessel (DLAV), (Figure 1 A timestamp 28:30, B). During anastomosis, dorsal movement of cells in ISVs decreased to averages of 2μm/h and 1μm/h for venous and arterial ISVs, respectively (Figure 1 B). Changes in velocity take place again at around 32 hpf (Figure 1 B). At this point in development, remodelling of ISVs begins and EC movement becomes significantly different between future aISVs and vISVs (Suppl Figure 2 A). For sprouts that remain arterial, the velocity continues to decrease, even reversing direction, averaging −0.6 μm/h from 34 to 38 hpf (Figure 1B). For sprouts that will remodel into veins, the orientation of movement remains the same, and velocity increases to an average to 3 μm/h (Figure 1 B). At 40 hpf aISV and vISV specification is complete, and EC velocity is reduced close to a zero-net velocity in all ISVs (Figure 1B, Supp. Figure 2). Overall, these results show that ECs decrease their velocity during DLAV formation (i.e. less dorsal movement of ECs). Later during remodelling, ECs from sprouts that remodel into venous ISVs increase their velocity (i.e. move dorsally), while ECs from sprouts that remain arterial (arterial ISVs) decrease their velocity and turn around (i.e. move ventrally).

### EC proliferation and exchange by directed migration control vessel diameter

To understand the role of differential EC movement in ISVs we followed the exchange of cells after DLAV anastomosis. To do so we measured how many ECs enter (immigration) or exit (emigration) the ISVs (Figure 1 C, schematic). After DLAV formation (after 31 hpf), ECs in the ISVs no longer move to the aorta, nor do ECs leave the dorsal aorta to enter aISVs (data not shown). In contrast, we find a tendency of cells to immigrate from the DLAV into aISVs. When measuring how often one or more cells enter the aISV, we observed that in 40% of all aISVs one cell enters these vessels and in 20% of all aISVs two cells enter (Figure 1 C, exchange with DLAV). No cells left the aISVs to contribute to the DLAV. Conversely, in vISV we find that cells emigrate towards the DLAV. We measured how often the emigration of cells towards DLAV happens and find a 56% fraction of vISVs with one cell and 20% with two cells emigrating (Figure 1 C, exchange with DLAV). No cells left the DLAV to contribute to the vISVs. However, since vISVs result from vessel remodelling, the connection to the PCV also constitutes a point of possible emigration or immigration of cells (Figure 1 D, pictogram). We find that the PCV only contributes with the emigration of ECs towards the vISVs, with 60% of vISVs gaining one cell and 32% gaining two cells (Figure 1 C, exchange with PCV). Thus the number of ECs in aISVs increases by immigration of cells from the DLAV, whereas vISV (Figure 1C exchange with DLAV) receive as many new cells from the PCV as they lose to the DLAV (Figure 1 C, exchange with PCV). Therefore, any apparent increase of ECs in vISVs needs to be achieved through a different process. We quantified the occurrence of cell proliferation by following the shape changes that cells go through during mitosis using the lifeact-GFP marker^17^ (cell rounding and increase of cortical F-actin) and counted how many times cell division occurs in venous and arterial ISV (Figure 1 A timestamp at 31:00 and 32:20). In aISVs, cell proliferation occurs once in a fraction of 50% and twice in a fraction of 10% of aISVs (Figure 1 E). In venous ISVs, the rate of proliferation is increased compared to aISVs, with a 46% fraction of two mitotic events in vISVs (Figure 1 E, Adapted Kolmogorov-Smirnov test, see methods, null test for frequency of mitosis in aISV vs vISV p<0.001). These results show that aISVs increase their number in ECs by a combined process of cell immigration, where cells enter the aISVs from the DLAV, and proliferation. In contrast, in vISVs an increase of ECs is ultimately dependent on cell proliferation (which happens more frequently compared to aISVs), since the movement of new cells immigrating from the PCV is balanced by the emigration of cells from vISVs towards the DLAV. Intriguingly, the differential movement of cells and rates of proliferation give rise to ISVs (both arterial and venous) that similarly contain between 3 and 4 cells (Figure 1 F, p>0.05). Moreover, these distinct processes accompany a steady and linear increase in vessel diameter as development progresses (Figure 1 G, diameter after remodelling: venous ISV ∼8 μm; arterial ISV ∼7 μm).

### Vascular morphogenesis in the zebrafish trunk requires the actin regulator WASp

Recent studies^22,23^ reported that EC polarity and directional movement are pre-specified in the developing zebrafish trunk ISV prior to their remodelling. Our current results support these findings and further identify a balance between oriented cell migration and vessel specific rates of proliferation during the establishment of a functional vessel morphology. To identify molecular mechanisms responsible for differential cell migration, we screened sorted ECs for the expression of known actin regulators by qPCR. From a collection of 40 actin regulators, the Wiskott–Aldrich Syndrome b (*wasb*) gene showed high levels of expression in ECs compared to non-endothelial cells at 24- and 48-hpf (Figure 2 and Supp Figure 3A). WASp was first isolated in lymphocytes^24^ and identified as an important regulator of the actin cytoskeleton of myeloid and lymphoid cells^25^. Endothelial expression or function of WASp (wasb) however has not been reported. To investigate a potential role of *wasb* in vascular remodelling, we knocked-out *wasb* with CRISPR-Cas9 (Supp Figure 3B). In F0 embryos (N=2) the trunk vasculature showed gross abnormalities. In all embryos, the aorta no longer ran parallel to the anterior-posterior axis of the embryo and instead showed bending and irregular diameter. Furthermore, the ISV’s stereotypical distribution across the trunk was lost and instead several aberrant connections formed between ISVs, the aorta and the PCV. Nonetheless, perfusion could still be observed (Figure 2 A and Supp Video 2). In previous studies focusing on the myeloid cell function of wasb, morpholinos against *wasb* specifically and efficiently knocked down *wasb*, producing the same myeloid cell phenotype reported for *wasb* mutants^26,27^. Therefore, we also made use of the same oligomers to study the role of *wasb* in vascular development of zebrafish embryos. *Wasb* morphants (*MO-wasb)* developed aberrant ISV connections (3-way connections that did not resolve as well as direct ISV – ISV connections, Figure 2C and Suppl. Figure 3C’) that persisted up to five days post-fertilization (Supp. Video 3). Furthermore, 54.5% of MO embryos showed ISVs with substantial abnormalities in vessel diameter (Figure 2C). These results suggest that *wasb* is required for correct vascular morphogenesis in the developing trunk vasculature.

**Figure 2.**
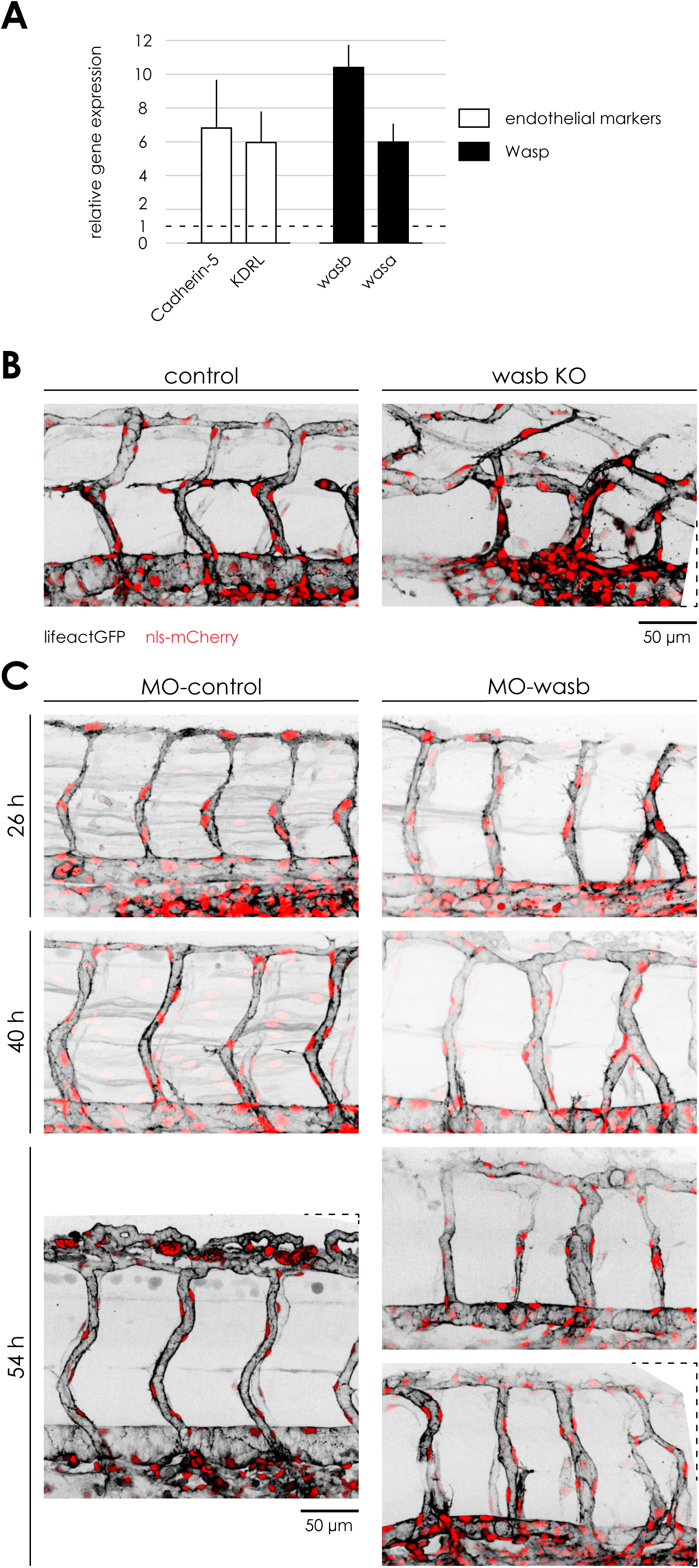
Vascular morphology is dependent of WASP. **A**, Reverse transcriptase PCR of FAC sorted ECs from Tg[*fliep:eGFP*] 24 hpf. Data are mean ±s.e.m. **B**. 44 hpf Tg[*fliep:nls-mCherry; fliep:lifeactGFP*] trunk vasculature of F0 embryo injected with Cas9 mRNA (control) and Cas9 mRNA plus guide mRNA (Was mutant). **C**, *MO-Ctr* and *MO-wasb* Tg[*fliep:nls-mCherry; fliep:lifeactGFP*] embryos. Top panel of *MO-wasb* shows large venous ISV with large diameter and arterial ISVs with reduced diameter. Lower panel an example of a vascular malformation.

### WASp controls endothelial cell migration

The Wiskott-Aldrich Syndrome protein (WASP) belongs to a family of nucleation promoting factors (NPF) known to bind to the Arp2/3 complex that drives the generation of branched actin for diverse cellular processes, including the formation of cellular protrusions required for cell migration^28^. Given that *MO-wasb* exhibit vascular morphology defects, we hypothesized that WASp may also play a role in the migration of ECs. To test this, we performed our quantitative morphodynamics analyses in *MO-wasb*.

Similar to control embryos, arterial sprouts from *MO-wasb* move with a positive velocity (i.e. dorsally) (Figure 3A, Supp Video 4). From 26 to 28 hpf ECs from *MO-Ctr* embryos move with an average velocity of 3.9 μm/h in aISVs and 5 μm/h in vISVs (Figure 1B). In *MO-wasb* these velocities were decreased to about half of the speed of controls, with an average of 2 μm/h and 2.5 μm/h for aISVs and vISVs, respectively (Figure 3B, Supp Figure 4). Like in *MO-Ctr* embryos from 28 until 30 hpf the DLAV formation is not impaired in *MO-wasb* (Figure 3A). However, while in *MO-Ctr* ECs decrease their velocity (1 μm/h for aISV and 3.5 μm/h for vISVs) in *MO-wasb* EC velocity is maintained (2 μm/h) for ECs in aISVs, and in vISVs there is an increase (4 μm/h) (Figure 3B). During vessel remodelling (after 31 hpf) in *MO-Ctr* embryos ECs show a differential behavior with ECs showing a decrease in velocity in aISVs and an increase in vISVs. In *MO-wasb* embryos EC velocity both in venous and arterial ISV velocity decreases (1 μm/h) and remains as such until 37 hpf, after which it reaches a net velocity of zero (Figure 3B). This result suggests that although WASp is not essential for anastomosis it appears to play a role in the regulation of ECs dynamic velocity.

**Figure 3.**
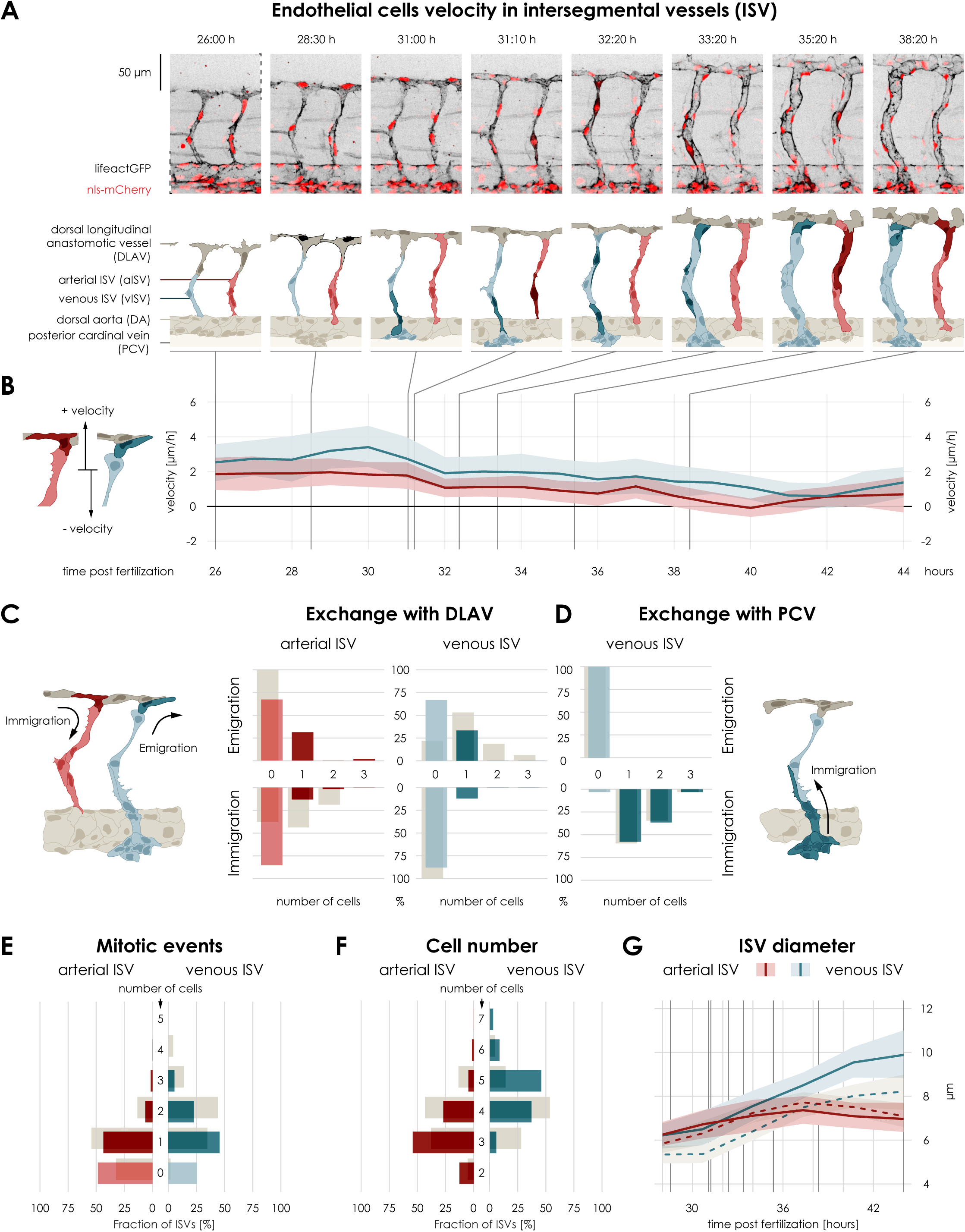
WASP is required for collective oriented cell migration. **A**, Montage of two ISVs sprouts from 26 hpf until 38 hpf from *MO-wasb* embryo. Top panel: Live imaging of Tg[*fliep:nls-mCherry; fliep:lifeactGFP*]. Black: F-actin, Red: nuclei. Bottom panel: Pictogram of live imaging; red ECs represent cells in aISVs, blue ECs represent cells in vISVs. Dark red ECs represent indicate cell division events (timepoint 31:10) and emigration of cells from aISV to DLAV (timepoints 35:20-38:20). Dark green cells indicate vascular remodelling (timepoint 31:00), cell division (timepoint 32:20, 33:20) and emigration of ECs towards the DLAV (timepoint 31:10, 33:20-38:20). Dark grey cells represent cell anastomosis (timepoint 28:30). **B**, Collective EC velocity (μm/h) across time post fertilization (hours). Red: velocity for ECs in aISVs, blue: velocity of ECs in vISV. Left pictogram demonstrates the referential for positive and negative velocity relative to dorsal and ventral movement. Data are mean ±CI (95%). Grey bars are values of *MO-Ctr*. **C**, Number of cells that exchange from ISVs with DLAV and fractions of ISVs that are affected. Grey bars are values of *MO-Ctr*. **D**, Number of cells that exchange from PCV with vISV and corresponding fractions of vISVs. Grey bars are values of *MO-Ctr*. **E**, Number of mitotic events in ISVs. Grey bars are values of *MO-Ctr*. **F**, Cell number in ISVs 44 hpf and distribution among ISVs. Grey bars are values of *MO-Ctr*. **G**, Diameter size (μm) across time post fertilization (hours) for *MO-Ctr* aISV (dotted redline), *MO-wasb* aISV (full redline), *MO-Ctr* vISV (dotted blue line) and *MO-wasb* vISV (full blue line). Black lines represent the equivalent time points in montage. Data are mean±CI (95%).

### WASp is required for differentially directed EC migration in arteries and veins

In search of the cellular processes leading to defective vascular patterning, we also quantified EC migration direction and cell exchange between ISVs and DLAV. In *MO-Ctr* embryos ECs from the DLAV immigrate into the aISV during remodelling (Figure 3C, immigration in arterial ISVs: 40% for one cell and 20% for two cells, grey bars). In *MO-wasb* embryos the immigration of cells into aISVs was substantially reduced (to one cell with a fraction of less than 20% of aISVs; Figure 3C, red bars). Moreover, unlike in MO-Ctr, in *MO-wasb* embryos some ECs moved from aISVs to the DLAV (with a frequency of 40% for one cell; Figure 3C, AKS test aISV exchange *MO*-Ctr vs. *MO-wasb* (p<0.001)). Cell proliferation however was unchanged (Figure 3D; AKS test p>0.05). Although the altered movement of cells from and into aISVs resulted in slightly more aISVs that harbor only 3 EC (Figure 3F, note: in *MO-Ctr* embryos vessels with 3 and 4 cells had equal frequencies. Grey bars), the overall number of ECs in the aISVs was not statistically different in MO-wasb compared to MO-Ctr (AKS test p>0.05) and the aISV diameter largely unaffected (Figure 3G, Suppl Figure 4).

In vISVs, however, altered cell movement patterns where even more pronounced, and ultimately responsible for the observed vascular patterning defects. Whereas in *MO-Ctr* vISV regularly contribute with one or more ECs that emigrate to the DLAV, this was strongly reduced in MO-wasb (compare Figure 1B and 1C to Figure 3C; AKS test p<0.001). Additionally, while in *MO-ctr* embryos ECs never immigrated from the DLAV to the vISVs, in *MO-wasb* this behavior did occur (with a 10% frequency; Figure 3C). Surprisingly, the contribution of cells from the PCV to vISVs was not different (Figure 3D AKS>0.05). Similar to aISVs, proliferation in vISVs was also unchanged (Figure 3E, AKS>0.05). The net effect of reduced emigration to the DLAV, maintained immigration from the PCV as well as the occasional immigration from the DLAV, was a significant increase in endothelial cell numbers in vISVs of MO-wasb (Figure 3 F), which paralleled the increased vessel diameter (Figure 3 G). Overall, this quantitative analysis suggests that WASp plays an important role in the coordination of directed movement of ECs during vessel remodelling.

### Arterial/venous wall shear stress is impaired in *MO-wasb*

During development, ECs are subjected to forces generated by blood flow in the form of wall shear stress (WSS). This force is determined by the flow rate, the blood viscosity, and the physical dimensions of blood vessels^29^. Given the alterations in vessel dimensions observed in embryos deficient in WASp expression, we asked how this altered network would impact flow and WSS within ISVs. We constructed an *in silico* stereotypical ISV network for each fish experiment, using measurements of nuclei displacement to instruct on the length of vessels and the measurements of vessel diameter changes to model the observed vessel morphology dynamics from 32 hpf to 44 hpf. This in silico model was then used to compute blood flow and WSS within aISVs and vISVs (Suppl. Figure 5A, B). When simulating flow in *MO*-Ctr embryos, WSS was initially higher in future vISVs. With further development however, aISVs emerged as the high WSS vessels (Suppl. Figure 5C). This shift in WSS mirrored the dynamic changes in vessel diameter, as aISVs were initially the larger diameter vessels at 32 hpf. As development progresses, vISV diameters increase (Figure 1 G), shifting the higher WSS to the aISVs. In *MO-wasb*, however, this WSS inversion never occurred and aISVs showed higher WSS from 32 hpf until 44 hpf (Suppl. Figure 5C). Interestingly, the WSS in aISVs steadily increased although their diameter remained constant. Closer examination revealed that this effect is the consequence of a global network effect driven by the enlarging vISVs (Suppl. Figure 5C and Figure 3G). These findings suggest that morphological changes in a particular set of vessels, like the increase of diameter in vISV observed in MO-wasb, have a global effect on WSS levels in otherwise normal looking vessels of the network.

### Wasp regulates endothelial actin organization

Given that Wasp stimulates actin nucleation, we asked how actin filaments are affected in ECs with decreased *wasb* expression. To address this, we imaged F-actin in ECs of Tg(*fliep:lifeactGFP)* embryos injected with *wasb* morpholino. At 26 hpf dorsal sprouts in *MO-wasb* showed a significant reduction of actin filaments (Figure 4A, compare upper panel at 26h PF control MO with lower panel *MO-wasb*). Although filopodia and lamellipodia were still visible in tip cells of these sprouts, the overall density of actin filaments in these structures was reduced (Figure 4A 26 hpf inset 2). This reduction was also visible at junctions between tip and stalk cells (Figure 4A 26 hpf panel 4). Actin bundles reminiscent of stress fibers were also lost (Figure 4A 26 hpf inset 3). Bundles of filamentous actin prominently mark the junctions of ECs (junctional actin)^17^ in the dorsal aorta of *MO-Ctr* injected embryos (Figure 4A 26h PF inset 1). Commencing remodelling, junctional actin in the dorsal aorta of *MO-wasb* was strongly reduced (Figure 4A 34h PF inset 1). ISVs showed similarly decreased F-actin along the cell cortex and cell junctions, being occasional labeled with actin puncta (Figure 4A 34 hpf inset 2 and 3). After vessel remodelling, loss of junctional F-actin in the aorta persisted (Figure 4A 48 hpf inset 1). In both venous and arterial ISVs junctional F-actin was decreased and we observed junction associated actin puncta, suggesting that the assembly of continuous actin filaments along the EC cell junctions is particularly dependent on WASp (Figure 4A 48h PF inset 2 and 3).

**Figure 4.**
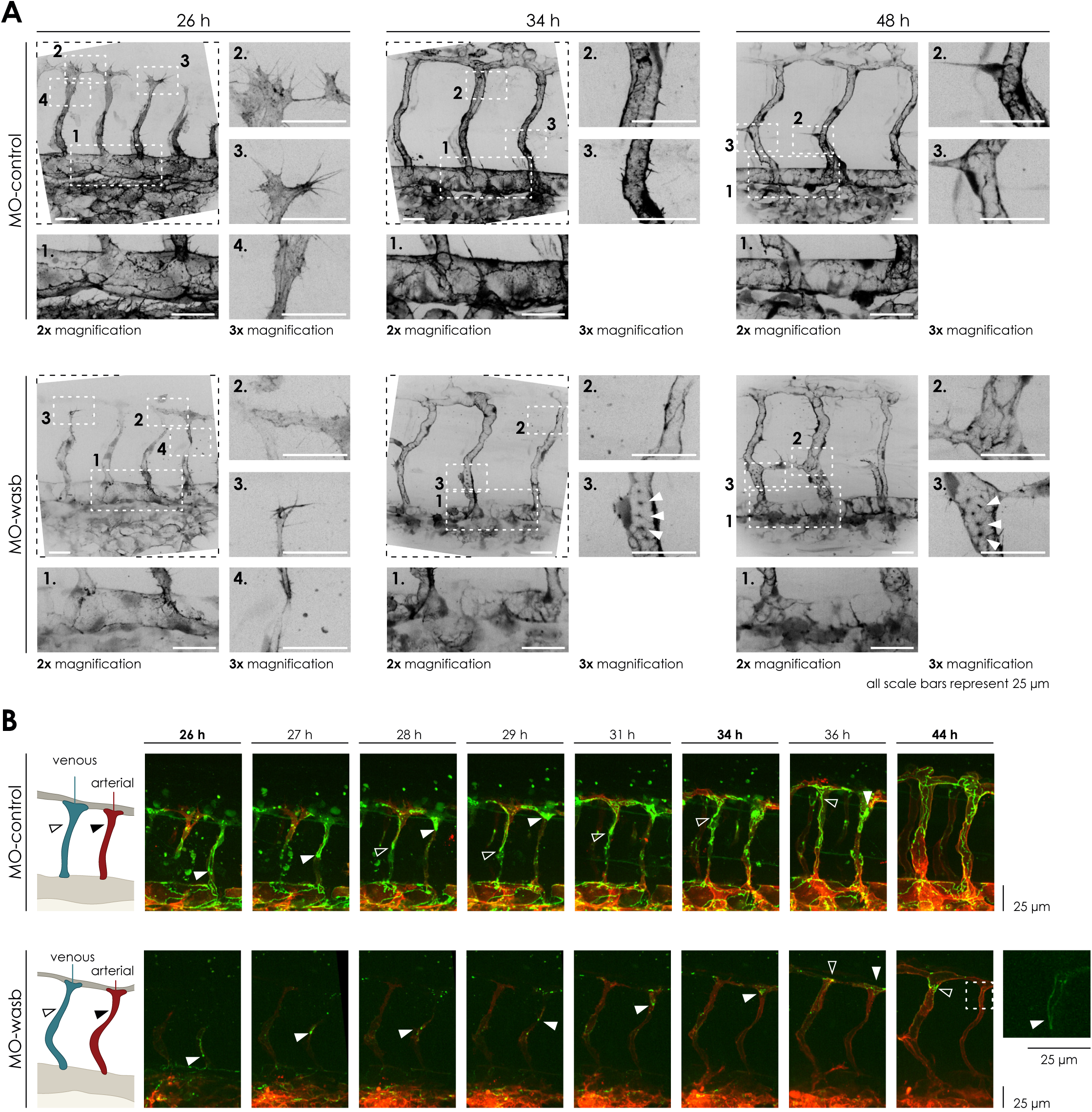
WAS is required for F-actin regulation and Junctional PECAM1 localization. **A**, Upper panel: Trunk vasculature of Tg[*fliep:lifeactGFP*] *MO-Ctr*. F-actin is mostly prominent at 26 hpf in sprouting ISV in filopodia (inset 3), cell-cell contacts in the initial steps of anastomosis (inset 2), in the cell cortex, as bundles resembling stress fibbers (inset 4). In the DA F-actin accumulates at cell junctions (inset 1). At 34 hpf and later at 48 hpf F-actin is enriched at cell junctions (junctional actin) in ISV (insets 2 and 3) at the DA (insets 1). Lower Panel: Trunk vasculature of Tg[*fliep:lifeactGFP*] *MO-wasb*. Overall F-actin in ECs is decreases, particularly in cell-cell contacts during anastomosis (inset 3) and heterogenous accumulation in the cell cortex and loss of stress fibber-like actin bundles (inset 4). Filopodia still show F-actin accumulation (inset 2). In the DA junctional F-actin is lost (Inset 1). At 34 hpf and later at 48 hpf junctional is heterogenous (insets 2) and in the form of puncta (insets 3). In the DA F-actin has heterogenous accumulation (insets 1). **B**, *MO-Ctr*: In green PECAM junctional localization in remodelling vessels labels migratory cells. White arrow head labels migratory stalk EC in aISv. Dorsal movement from 26-29 h. Diffuse PECAM localization at 34 hpf. Ventral movement at 36 hpf. Black arrowhead labels migratory stalk EC in vISV with dorsal movement from 28-36 hpf. *MO-wasb*: PECAM is lost or reduced in EC junctions. White arrowhead labels migratory stalk EC in aISVs with dorsal movement from 26 to 36 hpf. Inset labels second stalk cell with ventral to dorsal orientation. Black arrowhead labels migratory EC entering vISV from 36-44 hours. Red labels ECs membrane.

Sauter and colleagues proposed that the cell interface elongation required for angiogenic sprouting is dependent on junctional reorganization of cadherin-5 and actin polymerization^16^. This junctional elongation requires active actin polymerization and the cytoplasmic tail of cadherin-5 to interact with the actin-cytoskeleton. Treatment with Latrunculin B blocking general actin polymerization or knocking down the actin nucleator fmnl3 localizing to EC junctions^30^, both causes disrupted cell elongation, and endothelial cell rearrangement to establish longitudinal parallel junctions within the ISVs^16,30^. To investigate the impact of *wasb knockdown* on cell interface elongation and Cadherin-5 localization, we took advantage of a Cadherin labeling transgenic line *Tg*(*ve-cad:ve-cadTS*)^31^. Although VE-Cadherin-TS still localized to cell junctions in *Tg*(*ve-cad:ve-cadTS*), embryos injected with *wasb* morpholino, (Suppl. Figure 6), the most dorsal part of the ISVs lacked continuous junctional labelling. This result mirrors what is seen in embryos treated with Latrunculin B, indicating that WASp-dependent actin polymerization is required for junction elongation and remodelling.

### WASp is necessary for junctional localization of PECAM in ECs

To further characterize the junctional defects, we investigated PECAM-1 expression (also known as CD-31), a junctional protein with important roles in junctional integrity^32^ and cell migration^33–35^. To achieve this, we took advantage of *Tg[fli1a:pecam1-EGFP]*^*ncv27*^, that expresses a PECAM-EGFP fusion protein specifically in ECs^36^. In *Tg[fli1a:pecam1-EGFP]*^*ncv27*^ embryos, PECAM clearly labeled junctions of ECs in the aorta (Figure 4B). In sprouts and throughout vessel remodelling, PECAM junctional localization is diffuse (Figure 4B time interval from 26h-36h PF). However, by the end of ISV remodelling PECAM localization becomes junctional (Figure 4B time 44h PF). Interestingly, during vessel remodelling ECs with a noticeable junction accumulation of PECAM display migratory behavior and PECAM junctional distribution can be seen to accumulate at the cell rear as the cell displays oriented cell migration (Figure 4B white arrowhead for arterial ISV and black arrowhead for venous ISV). Thus, PECAM junctional accumulation informs on the orientation of the migratory cell behavior by accumulating at the cell rear (Figure 4B black arrowhead). Intriguingly, we observed when cells change their migration direction, PECAM localization becomes diffuse (Figure 4B control panel time point 29h-34h PF cell labeled with white arrowhead). However, just before the new direction is established, PECAM resumes its junctional localization (Figure 4B control panel time point 36h PF). In *Tg[fli1a:pecam1-EGFP]*^*ncv27*^ embryos injected with *wasb* morpholino, the junctional accumulation of PECAM in ECs of the aorta was reduced (Figure 4B *MO-wasb*). Also, in sprouts and throughout vessel remodelling, junctional accumulation of PECAM was severely impaired, with a few puncta accumulating at the rear end of migratory cells (Figure 4B *MO-wasb*, white arrowhead). Interestingly, as MO-wasb cells in aISVs fail to reverse their migratory direction, unlike in MO-Ctr, their residual PECAM expression never became diffuse (Figure 4B time point 44h PF see inset). These results suggest that WASp-mediated actin nucleation is required for repolarization of junctional PECAM involved in directional migration.

### WASp is required for oriented cell migration in human ECs

To determine whether this endothelial function of WASp is uniquely required during remodelling of blood vessels in zebrafish, we examined the role of human WASp in cultured Umbilical Vein Endothelial Cells (HUVECs). We studied the morphology, migration properties and accumulation of junctional proteins in HUVECs, treated with small interfering RNA (siRNA) against *was* (siWAS). In a confluent HUVEC monolayer treated with siRNA against *was* (siWAS), EC morphology changed from a characteristic cobblestone morphology to an elongated shape with cells organized into streams and swirls (Figure 5A). This collective arrangement of cells is highly similar to what was recently observed in HUVECs after the knock-down of YAP/TAZ, where cells adopt this very morphology as a consequence of reduced junctional remodelling and the inability to rearrange dynamically within the monolayer ^37^. Thus, WASp may be similarly required for ECs to shuffle between neighbors. To test the role of WASp in cell migration, we performed a wound closing assay^38^. HUVECs treated with siWAS were substantially less effective in closing the cell free area compared siRNA control cells (siCTR) (Figure 5B). 16h post scratch siWAS treated cells showed a 70% coverage of free space instead of 100% in siCTR treated cells (Figure 5C), suggesting defects in cell migration. To further investigate this hypothesis, we performed live imaging and tracked the nuclei displacement as a proxy for cell migration in the first row of cells. In siCTR experiments ECs exhibited a polarized movement towards the cell free area (Figure 5E) and a velocity of ∼25 μm/h (Figure 5 D). siWAS treated ECs instead migrated in all directions (Figure 6E), including parallel to the wound edge, and showed a significant decrease in velocity (Figure 5D, ∼10 μm/h). The polarization of follower cells (looking at the row of cells immediately behind the first row) was also affected (Supp Figure 7A) indicating a loss of collective coordination of polarity^39^. Therefore, like in the developing vasculature of *D*.*rerio* embryos, in human ECs WASp plays an important role in oriented cell migration.

**Figure 5.**
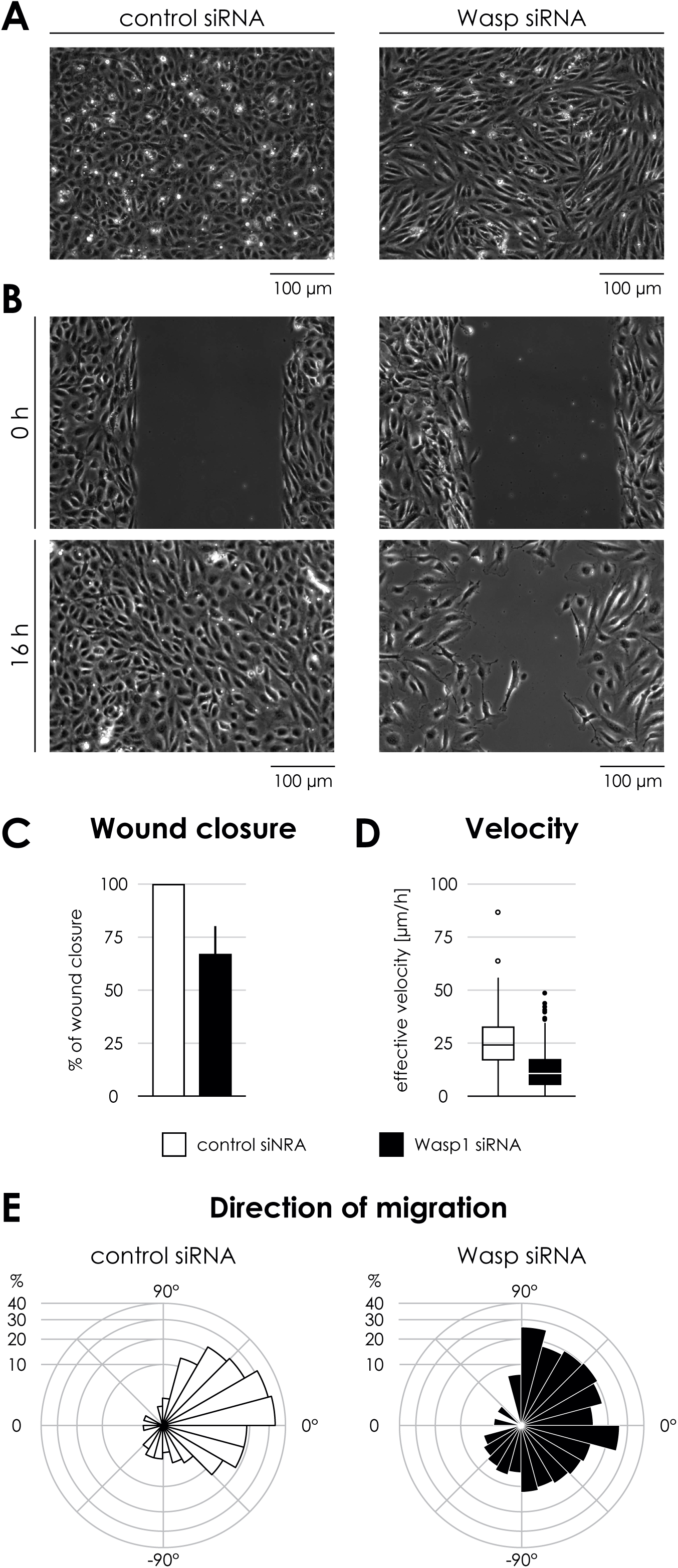
WASP controls migration and PECAM localization in HUVECs. **A**, Phase contrast images of Control and Was siRNA treated HUVECs in a confluent monolayer. **B**, Scratch wound assay for Ctr and Was siRNA treated HUVECs. 0 hours immediately after removing barrier to create a cell free space and 16 hrs later. **C**, Quantification of wound closure at 16 hrs. Data are mean ±SD of 3 independent experiments (6–7 biological replicates). **D**, Effective velocity of HUVECs during wound closure assay. Samples have a significant velocity difference, p-value< 0.0001 (Welch’s two sample t-test). **E**, Rosette graph showing the prevalent cell direction during wound closure in Ctr and Was siRNA treated cells. Comparison between graphs p-value<0.001 (Watson’s two-sample test of homogeneity).

**Figure 6.**
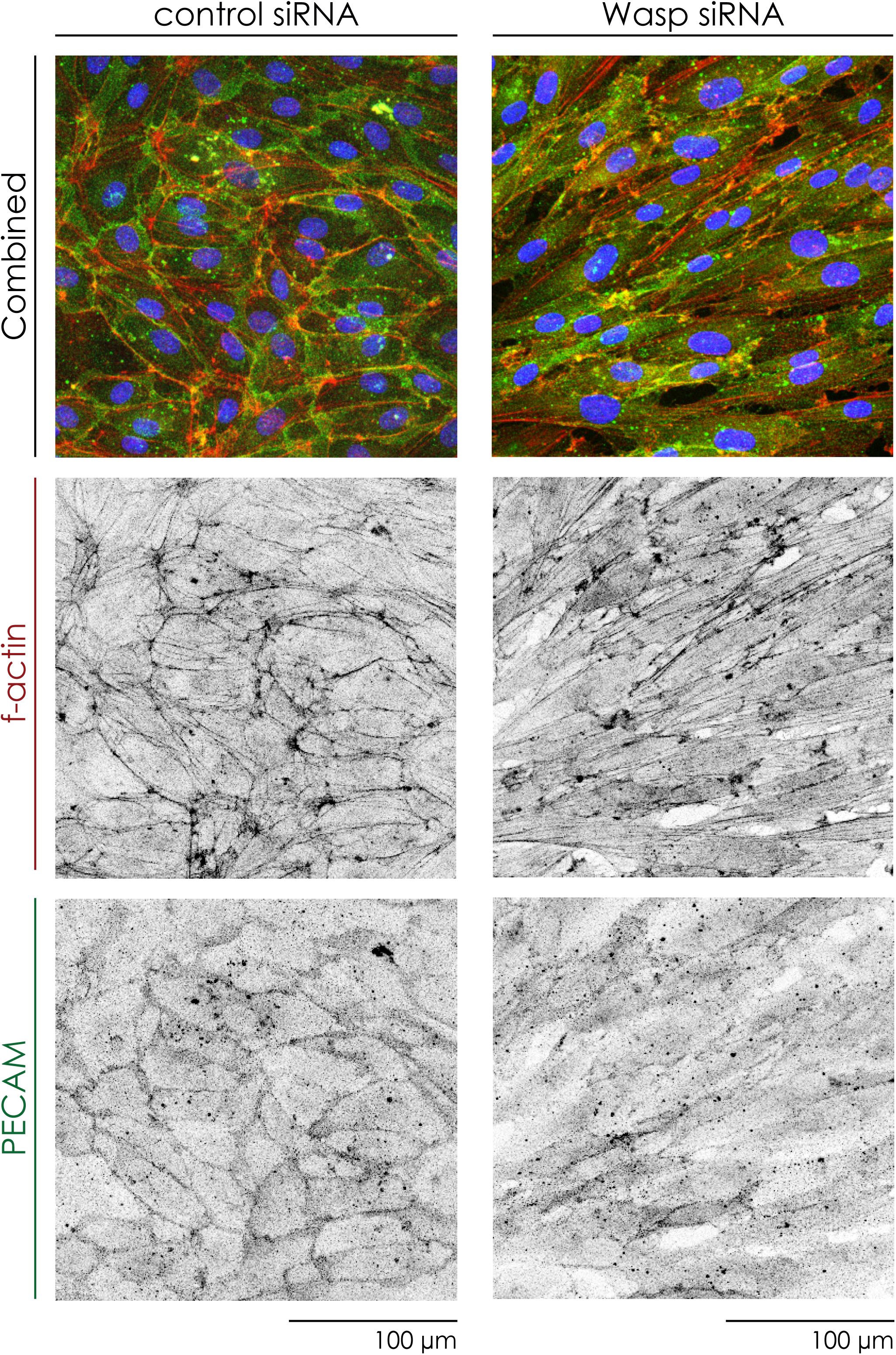
WASP is required for PECAM junctional localization. **A** HUVECs treated for Ctr and **B** Was siRNA stained for and f-actin (phalloidin, red in combined), PECAM1 (green in combined) and nuclei (DAPI, blue)

We further studied F-actin and PECAM organization to assess whether the role of wasb in zebrafish is conserved in HUVECs. siWAS treated HUVECs displayed an increase in actin bundles compared to siCTR cells (Figure 6 and Suppl. Figure 7B), suggesting that like Neural-WASP, WASP also prevents the accumulation of stress fibers^40^. Interestingly, we also observed a decrease in branched actin networks associated with adherens junctions (Suppl. Figure 7B) suggesting a decrease in junction associated intermediated lamellipodia (JAIL) formation, which are linked to cell migration^41^. Defects in JAIL formation would also agree with the observed decrease in cell migration velocity (Figure 6D). Finally, the loss of WASP in HUVECs greatly impaired junctional localization of PECAM in monolayers (Figure 7). Together, these results demonstrate an unappreciated role of WASP in the biology of zebrafish and human EC, critically regulating junctional actin and PECAM localization that are required for coordinated directional migration during vascular remodelling.

**Figure 7.**
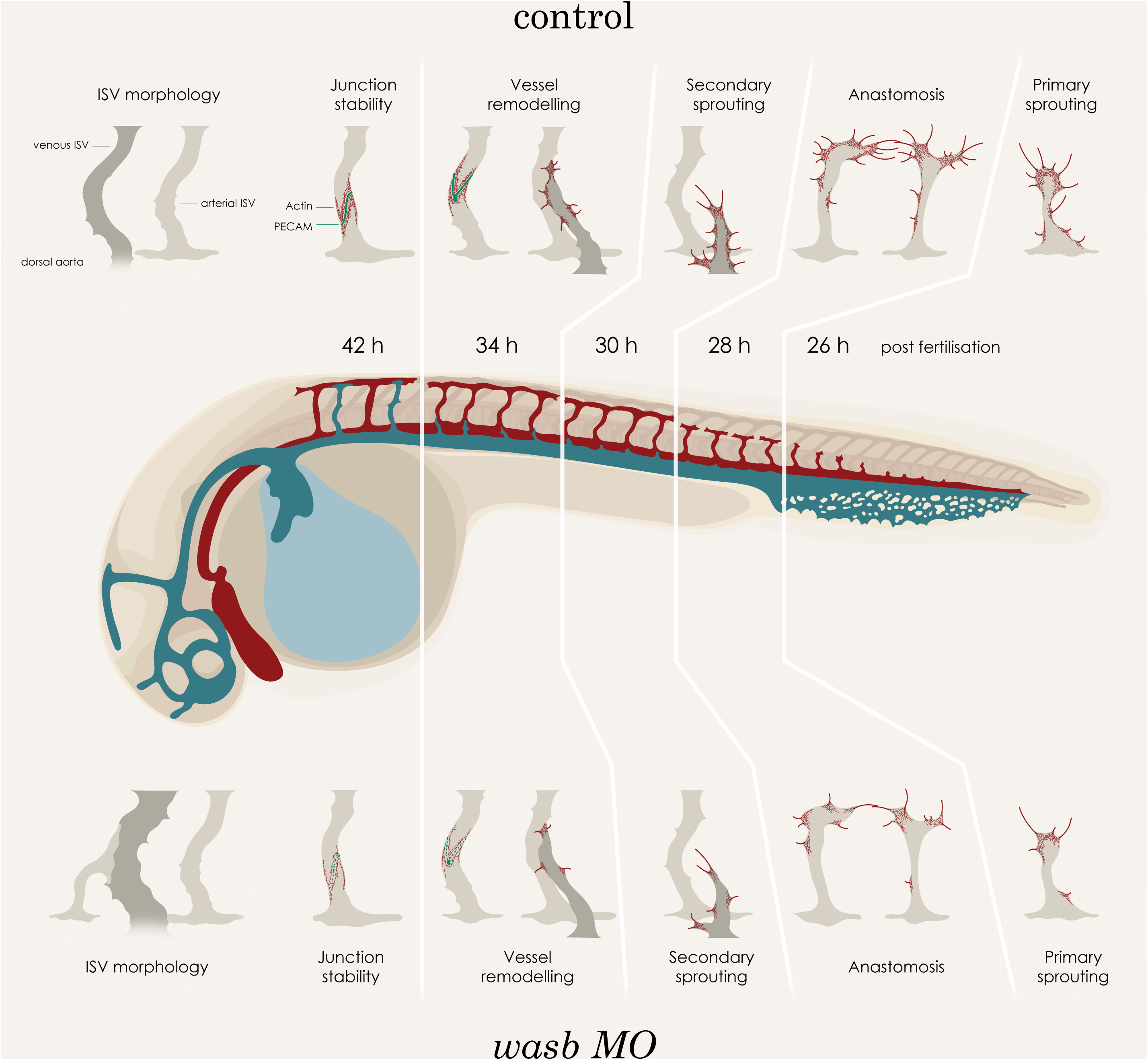
Illustration of zebrafish trunk vasculature development depicting morphological changes in intersegmental vessels and alterations in F-actin and junctional PECAM localization in key cellular process.

## Discussion

The present study provides two major insights into mechanisms of vascular patterning. Firstly, it predicts that diameter control in the highly organized vascular network in the zebrafish trunk is achieved through complex directional migration patterns that differ remarkably between future arteries and veins throughout the remodelling process. Accordingly, the regular numbers of endothelial cells in both arterial and venous ISVs is not primarily determined by proliferation within a given vessel segment, but rather by how many cells enter or exit from and to neighboring segments. We called this process immigration and emigration to capture the migratory behavior from the perspective of a given segment. The DLAV that arises from anastomosing primary sprouts^20^ here emerges not only as a crucial structure to provide lumenal continuity and thus a closed vascular loop to transport blood, but also as a conduit for endothelial exchange and rearrangements. This exchange of cells appears strikingly non-random, as movement from future venous ISVs into the DLAV, and from the DLAV into future arterial ISVs appeared to be the natural principle behavior. Quantifying all endothelial trajectories in the ISVs of around 20 embryos demonstrated an astonishing degree of consistency in this behavior, suggesting that it is tightly regulated.

Secondly, our present work identifies that the actin nucleation promoting factor WASp, previously believed to be expressed primarily in immune cells and associated to immune and blood deficiencies^25^, exerts a critical role in driving junctional F-actin assembly required for endothelial directional migration behavior. WASp is an integral player of the molecular machinery responsible for motility in immune cells, where *in vitro* and *in vivo* studies in cells of patients with the Wiskott-Aldrich syndrome and WASp-knockout mice revealed perturbed oriented migration^42,43^. In line with this, our results show that WASp function is conserved in ECs since impaired expression has consequences of disrupted differential migratory EC behavior, resulting in aberrant connections, dramatically increased and irregular diameter of veins, and a formation and persistence of arteriovenous shunts, phenotypically, and hemodynamically resembling human vascular malformations^44,45^. We propose that WASp-deficient ECs have impaired migration response that prevent the reorientation of directional motility required during vessel remodelling, a phenomenon analogous to the loss of navigation accuracy to stop and start to move again in leukocyte and macrophage WASp-deficient cells^27^. This principle of balanced directional migration provides a new conceptual framework to understand the emergence of vascular malformations, and suggests that proliferation alone is unlikely the exclusive driver of such pathologies.

Additionally, we show that WASP is required for the junctional accumulation of PECAM1. It’s unclear if this is a direct consequence of WASp-dependent actin polymerization and cytoskeletal rearrangement or of its activity as a scaffold protein^46^. Like Cadherin 5, PECAM1 has been shown to be involved in force transmission after mechanical stimuli^47^, assuming a “tug of war” of forces between cells to drive and coordinate movement, our current hypothesis is based on the function of WASp in the control of junctional actin remodelling and assembly of effective junctional proteins, responsible for signal propagation for oriented cell migration during vessel remodelling (Figure 7).

Overall, these results further support that the zebrafish trunk vasculature offers a unique and simplified window for the study of vessel establishment. For this reason, this model is particularly befitting to study diseases that involve the vascular system in a dynamic fashion, and to identify new mechanisms, cellular and molecular, involved in the formation of blood vessels. In this study we report the first analysis of WASp in a cellular context other than the immune system, challenging the view of restricted molecular pathways associated to F-actin regulation and migration of hematopoietic cells. Instead, we demonstrate that these mechanisms are at work in cells of hemangioblast origin, providing new avenues in the treatment of vascular diseases focusing on targets that are key players in the actin regulation and increasing the clinical differential for patients that suffer from Wiskott-Aldrich syndrome given the potential of vascular pathologies.

## Materials and methods

### Zebrafish husbandry and transgenic lines

Danio rerio were raised and staged as previously described^47^. The following transgenic lines were used: Tg[fli1a:pecam1-EGFP]ncv27 ^47^ (labels endothelial cell junctions), (Tg(kdr-l:ras-Cherry)s916 ^48^ (labels endothelial cell membrane), Tg(fli1ep:Lifeact-EGFP)^17^ (labels endothelial F-actin), Tg(fli1:NLS-mCherry)^49^ (labels all endothelial cell nuclei) and Tg(ve-cad:ve-cadTS)^31^ (labels Cadherin 5). For growing and breeding of transgenic lines, we complied with regulations of the animal ethics committees at MDC Berlin.

### Live imaging zebrafish embryos

Embryos were anaesthetized in 0.007% tricaine (MS-222, Sigma-Aldrich), mounted in a 35mm Sarstedt (Ref #82.1473) petri dish using 0.8% low melting point agarose (Sigma-Aldrich) and bathed in Danieau’s buffer containing 0.007% tricaine and 0.003% PTU. Time-lapse imaging was performed using upright 3i spinning-disc confocal using a Zeiss Plan-Apochromat 20×, 40× or 63×/1.0 NA water-dipping objective and heating chamber. Image processing was performed using Fiji software^50^.

### Morpholino Knock-down

Morpholinos against *wasb* were used as previously described^26,27^.

### Imaging of HUVECs

Time-lapse imaging was performed using a Carl Zeiss LSM780 inverted microscope with a Plan-Apochromat 20x/0.8 at 37°C under 5% CO2. Imaging of fixed samples were performed using a Carl Zeiss LSM700 upright microscope with a Plan-Apochromat 20x/0.8.

### Scratch wound assay

24 hr after siRNA transfection cells were re-plated into a scratch wound assay device (IBIDI). On the following day a cell free gap of 500 µm was created by removing the insert of the device. Live imaging was performed immediately after removing the insert. For cell coverage measurements images were taken immediately after removing the insert and after 16 hr using a Leica DMIL LED microscope equipped with a 10×/0.22 NA Ph1 objective and a CCD camera (DFC3000 G). The cell free area was measured in Fiji and used to calculate the percentage of wound closure at 16 hr.

### Immunofluorescence staining

For immunofluorescence in HUVECs, cells were grown in #1.5 coverslips coated with poly-lysine and gelatin 0.2%. At the end of the experiment cells were fixed in 4% PFA for 10 min, permeabilised in 0.3% Triton-X100 in blocking buffer for 5 min and blocked in 1% BSA 20 mM Glycine in PBS for 30 min. Primary and secondary antibodies were incubated for 2 and 1 hr, respectively, in blocking buffer. Nuclei labelling was performed by incubating cells with DAPI for 5 min (Life technologies D1306) and Alexa Fluor 568 Phalloidin (1/600, ThermoFisher A12380). Primary antibodies used were as follows: Human VE-Cadherin (1/100, R&D AF938), human PECAM1 (1/200, Abcam ab76533).

### Segmentation and tracking of endothelial nuclei

Segmentation and tracking of nuclei from live imaging in zebrafish and cultured HUVECs was performed with Imaris Image Tracking Package (Bitplane). Tracking data was formatted according to community standards for open cell migration data^51^. In zebrafish the cell tracking data was oriented into three-dimensional space such that the aorta is parallel to the x-axis and the DLAV in y’s positive half section. In this way, the y-component of the transformed trajectories serve as a read-out for the ventral to dorsal positions of the nuclei. In order to determine the orientation of the aorta we used positional data of all nuclei in the aorta over the whole time period and performed linear regression The tracking was subsequently rotated and translated in three-dimensional space (see SI text for details). We extracted ventral to dorsal velocities over the developmental process, by calculating the signed displacement in y-direction over time windows of 2 h. This process was repeated over the whole time period from 26 h to 44 h in 1 h time steps for all cell tracks. In HUVECs cell trajectory data was aligned such that the y-axis followed the outer edge of the scratch-wound, while the cell-free space is on the positive-x half plane. We calculated the effective speed as the quotient of the distance from start to end point and the tracking time. Furthermore, we determined the directionality of each trajectory from the vector pointing from start to end point and calculated the angle with the y-axis. The data that support the findings of this study are available from the corresponding author upon reasonable request.

### Crispr-Cas9 Knock out

Guide RNA design was performed making use of ZiFit Targeter software package. RNA guide with sequence TAGGAATGGAGTCTCCAGCATAC was used to target exon 2 of zebrafish wasb. Cas9 mRNA and guide RNA was injected at one cell stage embryos of Tg(fli1ep:Lifeact-EGFP);Tg(fli1:NLS-mCherry).

### Cell disruption for FAC sorting

Dechorination of zebrafish embryos was performed by gently pipetting embryos up and down in a solution of Danieau’s buffer containing pronase. After dechorination embryos were transferred to a Calcium free Ringer solution (116mM NaCL, 2.9 mM KCl, 5mM HEPES pH 7.2). Yolk was removed by gently pipetting. Centrifuged (720 rcf, Eppendorf 5804 R) for 5’ at 4°C to remove supernatant. Dissociation was performed by re-suspension in liberase solution (0.8 mg/ml liberase in DPBS) at 28.5°C for 10’. To stop dissociation samples were placed on ice and added to a solution with 1-2% CaCl2 and 5-10% FBS. Finally, centrifuge for 5’ at 4°C, discard supernatant and resuspend DPBS with 2mM of EDTA. FAC sorting was performed in a BD Aria sorter at the MDC flow cytometry facility. Sorted cells were kept at −80°C in Trizol.

### RNA extraction and real-time quantitative RT-PCR

RNA extraction was performed making use of Direct-zolTM RNA MicroPrep (Zymo Research, R2060, R2061, R2062, & R2063). First strand cDNA synthesis was performed with Thermo Scientific RevertAid Reverse Transcriptase kit (Thermo Scientific, EP0441) with random hexamer primers for cells extract from zebrafish embryos and RNeasy plus Mini Kit (Qiagen, 74034) for HUVEC samples. SYBR Green quantitative real-time quantitative PCR was performed following SG qPCR Master mix protocol and reagents from Roboklon (EURx Roboklon, E0402-01) for zebrafish samples Primers used for quantitative RT PCR were as published^27^. For HUVEC samples real-time quantitative PCR was performed using TaqMan reagents (Applied Biosystems).

### *In silico* idealized ISV network

We constructed an idealised model of flow within the ISV network for each zebrafish experiment using summary statistics in order to estimate flow/WSS experienced by ECs. Each vessel network was represented as a graph (i.e., a collection of nodes and edges) obtained by linear regression of the cell trajectory data. Each network consisted of 30 ISVs with an alternating pattern of aISV - vISV. The ISVs were positioned within the network according to mean ISV spacing and ISV length measured from the experiment. Each edge was also assigned a diameter function over time based on mean diameter measurements of each vessel type (dorsal aorta, DLAV, aISV, or vISV). The distinction of aISV or vISV came from end-point analysis, i.e. designating the vessel by which vessel the ISV was connected to at the end of the experiment (either artery or vein).

Boundary nodes were defined at the inlet and outlet of the dorsal aorta, as well as at each of the terminal ends of the vISVs. Due to difficultly in imaging and distinguishing the pectoral cardinal vein from the dorsal aorta, the vein wasn’t included in the network and instead was treated as a flow sink (*P*_PCV_= 0 Pa). Pressure at the inlet was estimated from measurements from with the dorsal aorta of adult zebrafish^52^ (*P*_DA,in_= 201.3 Pa). A pressure gradient was prescribed along the aorta in order to maintain forward flow through the vessel over time. We estimated that this gradient dissipated the inlet pressure over a distance over the length of whole fish, *L*_fish_= 2 mm, and applied this gradient to the fraction of the aorta included in our flow model,

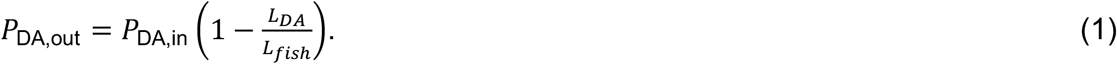

The flow conductance (i.e., the inverse of flow resistance) of each edge was calculated as

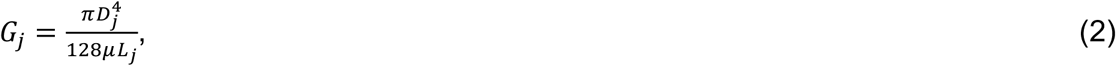

with *D*_*j*_ and *L*_*j*_ as the diameter and length of edge *j* at time point *t*, and *μ* the dynamic viscosity of blood (*μ* = 0.0035 kg/m/s). We use a flow balance equation at each node to assemble the linear system of equations for the network, which we can use to solve for the unknown nodal pressures and therefore flow based on vessel conductivity,

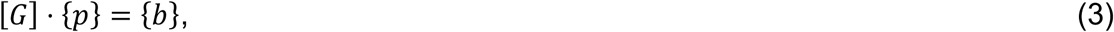

where {*p*} is the array of unknown nodal pressures, [*G*] is network conductivity matrix assembled with edge connectivity and conductivity, and {*b*} is the solution array which enforces the flow balance and contains information on the pressure boundary conditions (see^53^ for a detailed description). We solved this system of equations for each time point, each time updating [*G*] based on the measurements of vessel diameter obtained from the experiments. Edges with no vessel diameter measurement (i.e., *D*_*j*_= 0) had their conductance set to an infinitesimally small value (∼ 10^−30^) as to keep [*G*] invertible.

We used MATLAB’s *mldivide* operator (Release 2018b, The MathWorks Inc., Natick, MA, USA) to solve the linear system of equations for the nodal pressures, with which we can calculate the pressure difference across each vessel segment and thus the flow, *Q*, and wall shear stress, *τ*, for each edge,

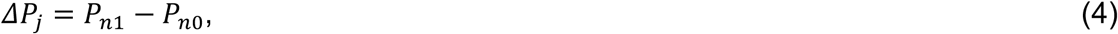

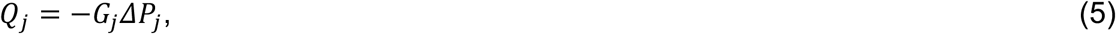

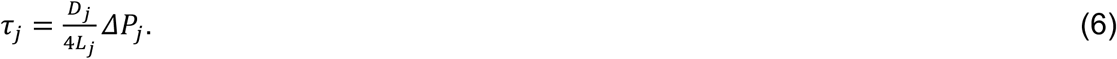

The data that support the findings of this study are available from the corresponding author upon reasonable request.

### HUVEC gene silencing

For knockdown experiments, HUVECs were transfected with SMARTpool: siGENOME siRNAs purchased from Dharmacon (was M-028294-02-0005, non-targeting siRNA Pool 1 #D001206-13-05). Briefly, subconfluent (70–80%) HUVECs were transfected with 25 nM siRNA using Dharmafect 1 transfection reagent following the protocol from the manufacturer; transfection media was removed after 24 hr and experiments were routinely performed on the second day after transfection.

### Computational analysis pipeline

For computational analysis we used the python-based workflow manager *kedro*, the code can be accessed on github https://github.com/wgiese/zebrafish_ec_migration. For all zebrafish and HUVECs experiments we created two key files, which contain meta data on the experimental setup as well as references to cell tracking data. Based on the key files our generic data pipeline allows to link data sets comprising cell trajectories, vessel diameters, vessel cell number, cell mitosis and inter-vessel cell migration (see Supp. Figure 1). The MATLAB code is available at https://github.com/ltedgar-ed/flow_idealised_ISV_network.

### Statistical analysis

For statistical analysis of cell number, cell divisions and inter-vessel migration distributions (see Supplemental Figures 2 and 4), we employed a modified version of the Kolmogorov-Smirnov test for discrete distributions by using the r-package dgof (insert ref from https://cran.r-project.org/web/packages/dgof/citation.html). The analysis of the HUVEC orientation data was performed based on the r-packages CircStats and circular, which provide statistical analysis of periodic data, including mean, standard deviation and Watson’s U2 test for circular distributions.

## Bibliography

1. Eilken, H. M. & Adams, R. H. Dynamics of endothelial cell behavior in sprouting angiogenesis. Curr. Opin. Cell Biol. 22, 617–625 (2010).

2. Herbert, S. P. & Stainier, D. Y. R. Molecular control of endothelial cell behaviour during blood vessel morphogenesis. Nat. Rev. Mol. Cell Biol. 12, 551–564 (2011).

3. Potente, M., Gerhardt, H. & Carmeliet, P. Basic and therapeutic aspects of angiogenesis. Cell vol. 146 873–887 (2011).

4. Betz, C., Lenard, A., Belting, H. G. & Affolter, M. Cell behaviors and dynamics during angiogenesis. Development (Cambridge) vol. 143 2249–2260 (2016).

5. Wacker, A. & Gerhardt, H. Endothelial development taking shape. Current Opinion in Cell Biology vol. 23 676–685 (2011).

6. Franco, C. A. et al. Dynamic endothelial cell rearrangements drive developmental vessel regression. PLoS Biol 13, e1002125 (2015).

7. Udan, R. S., Vadakkan, T. J. & Dickinson, M. E. Dynamic responses of endothelial cells to changes in blood flow during vascular remodeling of the mouse yolk sac. Development 140, 4041–4050 (2013).

8. Kochhan, E. et al. Blood Flow Changes Coincide with Cellular Rearrangements during Blood Vessel Pruning in Zebrafish Embryos. PLoS One 8, e75060 (2013).

9. Chen, Q. et al. Haemodynamics-Driven Developmental Pruning of Brain Vasculature in Zebrafish. PLoS Biol. 10, e1001374 (2012).

10. Michaelis, U. R. Mechanisms of endothelial cell migration. doi: 10.1007/s00018-014-1678-0.

11. Santos-Oliveira, P. et al. The Force at the Tip - Modelling Tension and Proliferation in Sprouting Angiogenesis. PLoS Comput. Biol. 11, e1004436 (2015).

12. Norton, K. A. & Popel, A. S. Effects of endothelial cell proliferation and migration rates in a computational model of sprouting angiogenesis. Sci. Rep. 6, 1–10 (2016).

13. Svitkina, T. The actin cytoskeleton and actin-based motility. Cold Spring Harb. Perspect. Biol. 10, (2018).

14. Heng, Y. W. & Koh, C. G. Actin cytoskeleton dynamics and the cell division cycle. International Journal of Biochemistry and Cell Biology vol. 42 1622–1633 (2010).

15. Angulo-Urarte, A. et al. Endothelial cell rearrangements during vascular patterning require PI3-kinase-mediated inhibition of actomyosin contractility. Nat. Commun. 9, 1–16 (2018).

16. Sauteur, L. et al. Cdh5/VE-cadherin promotes endothelial cell interface elongation via cortical actin polymerization during angiogenic sprouting. Cell Rep. 9, 504–513 (2014).

17. Phng, L. K., Stanchi, F. & Gerhardt, H. Filopodia are dispensable for endothelial tip cell guidance. Development 140, 4031–4040 (2013).

18. Franco, C. A. et al. Non-canonical Wnt signalling modulates the endothelial shear stress flow sensor in vascular remodelling. Elife 5, (2016).

19. Rørth, P. Collective Cell Migration. Annu. Rev. Cell Dev. Biol. 25, 407–429 (2009).

20. Hogan, B. M. & Schulte-Merker, S. Developmental Cell How to Plumb a Pisces: Understanding Vascular Development and Disease Using Zebrafish Embryos. (2017) doi: 10.1016/j.devcel.2017.08.015.

21. Heckel, E. et al. Oscillatory flow modulates mechanosensitive klf2a expression through trpv4 and trpp2 during heart valve development. Curr. Biol. 25, 1354–1361 (2015).

22. Weijts, B. et al. Blood flow-induced Notch activation and endothelial migration enable vascular remodeling in zebrafish embryos. Nat. Commun. 9, (2018).

23. Geudens, I. et al. Artery-vein specification in the zebrafish trunk is pre-patterned by heterogeneous Notch activity and balanced by flow-mediated fine-tuning. Dev. 146, 1–13 (2019).

24. Derry, J. M., Ochs, H. D. & Francke, U. Isolation of a novel gene mutated in Wiskott-Aldrich syndrome. Cell 79, 635–644 (1994).

25. Thrasher, A. J. & Burns, S. O. WASP: A key immunological multitasker. Nat. Rev. Immunol. 10, 182–192 (2010).

26. Jones, R. A. et al. Modelling of human Wiskott-Aldrich syndrome protein mutants in zebrafish larvae using in vivo live imaging. J. Cell Sci. 126, 4077–4084 (2013).

27. Cvejic, A. et al. Analysis of WASp function during the wound inflammatory response - Live-imaging studies in zebrafish larvae. J. Cell Sci. 121, 3196–3206 (2008).

28. Burns, S., Thrasher, A. J., Blundell, M. P., Machesky, L. & Jones, G. E. Configuration of human dendritic cell cytoskeleton by Rho GTPases, the WAS protein, and differentiation. Blood 98, 1142–1149 (2001).

29. Ballermann, B. J., Dardik, A., Eng, E. & Liu, A. Shear stress and the endothelium. in Kidney International, Supplement vol. 54 S100–S108 (Elsevier, 1998).

30. Phng, L. K. et al. Formin-mediated actin polymerization at endothelial junctions is required for vessel lumen formation and stabilization. Dev Cell 32, 123–132 (2015).

31. Lagendijk, A. K. et al. Live imaging molecular changes in junctional tension upon VE-cadherin in zebrafish. Nat. Commun. 8, (2017).

32. Privratsky, J. R. et al. Relative contribution of PECAM-1 adhesion and signaling to the maintenance of vascular integrity. J. Cell Sci. 124, 1477–1485 (2011).

33. Cao, G. et al. Involvement of human PECAM-1 in angiogenesis and in vitro endothelial cell migration. Am. J. Physiol. - Cell Physiol. 282, (2002).

34. Zhu, J. X., Cao, G., Williams, J. T. & DeLisser, H. M. SHP-2 phosphatase activity is required for PECAM-1-dependent cell motility. Am. J. Physiol. - Cell Physiol. 299, 854–865 (2010).

35. O’Brien, C. D., Cao, G., Makrigiannakis, A. & DeLisser, H. M. Role of immunoreceptor tyrosine-based inhibitory motifs of PECAM-1 in PECAM-1-dependent cell migration. Am. J. Physiol. - Cell Physiol. 287, 1103–1113 (2004).

36. Ando, K. et al. Clarification of mural cell coverage of vascular endothelial cells by live imaging of zebrafish. Dev. 143, 1328–1339 (2016).

37. Neto, F. et al. YAP and TAZ regulate adherens junction dynamics and endothelial cell distribution during vascular development. Elife 7, (2018).

38. Jonkman, J. E. N. et al. An introduction to the wound healing assay using live-cell microscopy. Cell Adhes. Migr. 8, 440–451 (2014).

39. Carvalho, J. R. et al. Non-canonical Wnt signaling regulates junctional mechanocoupling during angiogenic collective cell migration. Elife 8, (2019).

40. Rajput, C. et al. Neural Wiskott-Aldrich syndrome protein (N-WASP)-mediated p120-catenin interaction with Arp2-Actin complex stabilizes endothelial adherens junctions. J Biol Chem 288, 4241–4250 (2013).

41. Cao, J. et al. Polarized actin and VE-cadherin dynamics regulate junctional remodelling and cell migration during sprouting angiogenesis. Nat. Commun. 8, 1–20 (2017).

42. Snapper, S. B. et al. WASP deficiency leads to global defects of directed leukocyte migration in vitro and in vivo. J. Leukoc. Biol. 77, 993–998 (2005).

43. Zicha, D. et al. Chemotaxis of macrophages is abolished in the Wiskott-Aldrich syndrome. Br. J. Haematol. 101, 659–665 (1998).

44. Folkman, J. & D’Amore, P. A. Blood vessel formation: What is its molecular basis? Cell vol. 87 1153–1155 (1996).

45. Cox, J. A., Bartlett, E. & Lee, E. I. Vascular malformations: A review. Semin. Plast. Surg. 28, 58–63 (2014).

46. Charras, G. & Yap, A. S. Tensile Forces and Mechanotransduction at Cell–Cell Junctions. Current Biology vol. 28 R445–R457 (2018).

47. Conway, D. E. & Schwartz, M. A. Mechanotransduction of shear stress occurs through changes in VE-cadherin and PECAM-1 tension: Implications for cell migration. Cell Adh. Migr. 9, 335–339 (2015).

48. Hogan, B. M. et al. Ccbe1 is required for embryonic lymphangiogenesis and venous sprouting. Nat. Genet. 41, 396–398 (2009).

49. Heckel, E. et al. Oscillatory flow modulates mechanosensitive klf2a expression through trpv4 and trpp2 during heart valve development. Curr. Biol. 25, 1354–1361 (2015).

50. Schindelin, J. et al. Fiji: An open-source platform for biological-image analysis. Nature Methods vol. 9 676–682 (2012).

51. Gonzalez-Beltran, A. N. et al. Community standards for open cell migration data. Gigascience 9, 1–11 (2020).

52. Hu, N., Joseph Yost, H. & Clark, E. B. Cardiac morphology and blood pressure in the adult zebrafish. Anat. Rec. 264, 1–12 (2001).

53. Pries, A. R., Secomb, T. W., Gaehtgens, P. & Gross, J. F. Blood flow in microvascular networks. Experiments and simulation. Circ. Res. 67, 826–834 (1990).

