## Supplementary figure legend for "Wasp controls oriented migration of endothelial cells to achieve functional vascular patterning"

Supplementary Figures

Supplementary Figure 1: Data management and analysis pipeline: **A**, Cell migration data, morphological data on the vessel networks as well as data on cell number and mitosis events is linked via a key file. **B**, The data pipeline manager kedro was used to process and analyse the data. Each raw, intermediate, or further processed data set is registered in a data catalogue. A data set is typically a set of plots or spreadsheets. The workflow was visualized as a data processing graph, where each square corresponds to a python function and each circle corresponds to a data set. **C,** Examples for intermediate, primary, and reporting data. **D**, Data flow and folder structure from raw to reporting data.

Supplementary Figure 2: Statistical test and sample size: **A**, Welch's t-test for means of ECs velocity in aISV and vISV across development. Faint velocity graph (same as Figure 1B) to compare Welch's test with means of velocity. **B**, Distribution of the total number of fish (Grey bars) and tracks made by nuclei displacement (yellow bars) across development used to calculate EC velocity. **C-D**, number of fishes analysed per ISV across development.

Supplementary Figure 3: Vascular morphology is dependent of WASP. **A**, Reverse transcriptase PCR of FAC sorted ECs from Tg[*fliep:eGFP*] 48 hours PF. **B**, Small deletion in Exon 2 of *wasb* gene in Crispr-Cas9 *D. rerio* embryos. **C**, Quantification of vascular phenotype in Ctr and Was MO embryos. **C'**, Examples of trunk vasculature in embryos with no, moderate and strong phenotype

Supplementary Figure 4: Statistical test and sample size: **A**, Welch's t-test for means of ECs velocity in aISV and vISV across development for Was MO embryos. Faint velocity graph (same as Figure 3B) to compare Welch's test with means of velocity. **B**, Distribution of the total number of fish (Grey bars) and tracks made by nuclei displacement (yellow bars) across development used to calculate EC velocity. **C-D**, number of fishes analysed per ISV across development.

Supplementary Figure 5: VE-Cadherin is present in cell junctions despite problems in cell intercaltion. Ctr and Was MO trunk vasculature of *Tg*(*ve-cad:ve-cadTS*) at 26 hours and 32 hours PF. White bars show dorsal junctional discontinuity.

Supplementary Figure 6. WSS progression in ISV is affected in WASP MO embryos. **A**, Live imaging of whole Tg[(fli1ep:Lifeact-EGFP); Tg(fli1:NLS-mCherry)] with F-actin label in black and EC nuclei in red. ISV Sections 1-3 mark the region of diameter measurements across development. **B**, *In silico* model made of 30 ISV (further details see methods section). Black bar with arrow highlights the region of 3 aISV and 3 vISV used to follow WSS inference across time depicted in C. **C,** WSS inference of aISV and vISV in *MO-Ctr* and *MO-wasb* conditions.

Supplementary Figure 7: **A.** Nuclei tracking of HUVECs treated with SiCTR and SiWASP at the very edge of wound (ROW1) the line of cells just before (ROW2). Rosette graph showing the prevalent cell direction during wound closure in Ctr and Was SiRNA treated cells. Comparison between graphs p-value<0,001 (Watson Two-Sample test for homogeneity). **B** VE-Cadherin, F-actin and nuclei in SiCTR and SiWAS HUVECs in a monolayer. Green:VE-Cadherin, Red: F-actin (Phaloidin), Blue: DAPI
