## Supplementary movie legend for "Wasp controls oriented migration of endothelial cells to achieve functional vascular patterning"

Supplementary movies

Supplementary movie 1: Time-lapse of 26h PF trunk vasculature of *MO-Ctr* Tg[(fli1ep:Lifeact-EGFP); Tg(fli1:NLS-mCherry)] embryo.

Supplementary movie 2: Time-lapse of 44h PF trunk vasculature of *was* Crispr/Cas KO Tg[(fli1ep:Lifeact-EGFP); Tg(fli1:NLS-mCherry)] embryo.

Supplementary movie 3: Time-lapse of 5 days PF trunk vasculature of *MO-wasb* KO Tg[(fli1ep:Lifeact-EGFP); Tg(fli1:NLS-mCherry)] embryo.

Supplementary movie 4: Time-lapse of 26h PF trunk vasculature of *MO-wasb* Tg[(fli1ep:Lifeact-EGFP); Tg(fli1:NLS-mCherry)] embryo.
