## Supplementary figures and images for "Wasp controls oriented migration of endothelial cells to achieve functional vascular patterning"

### Supplementary Figures

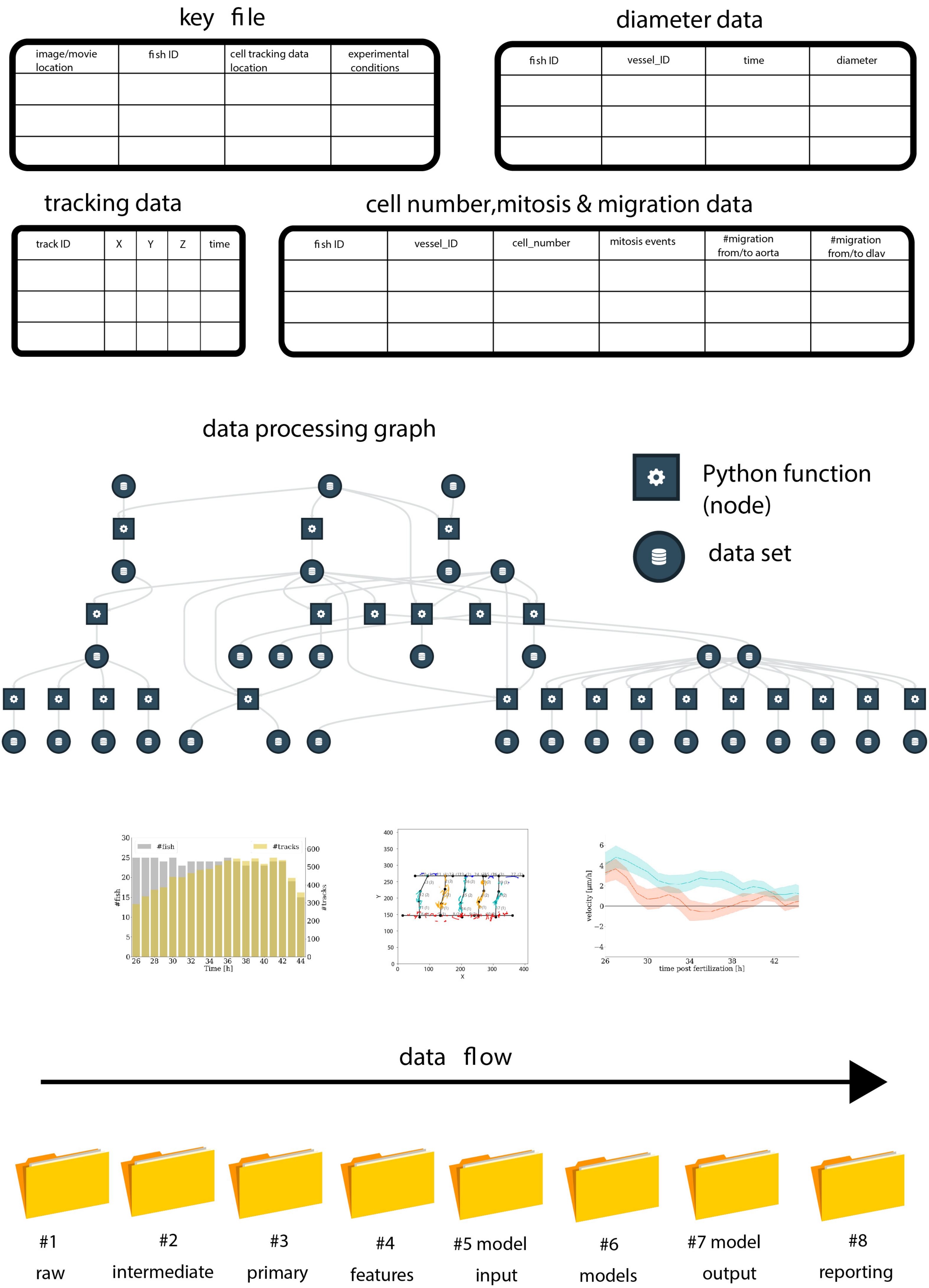

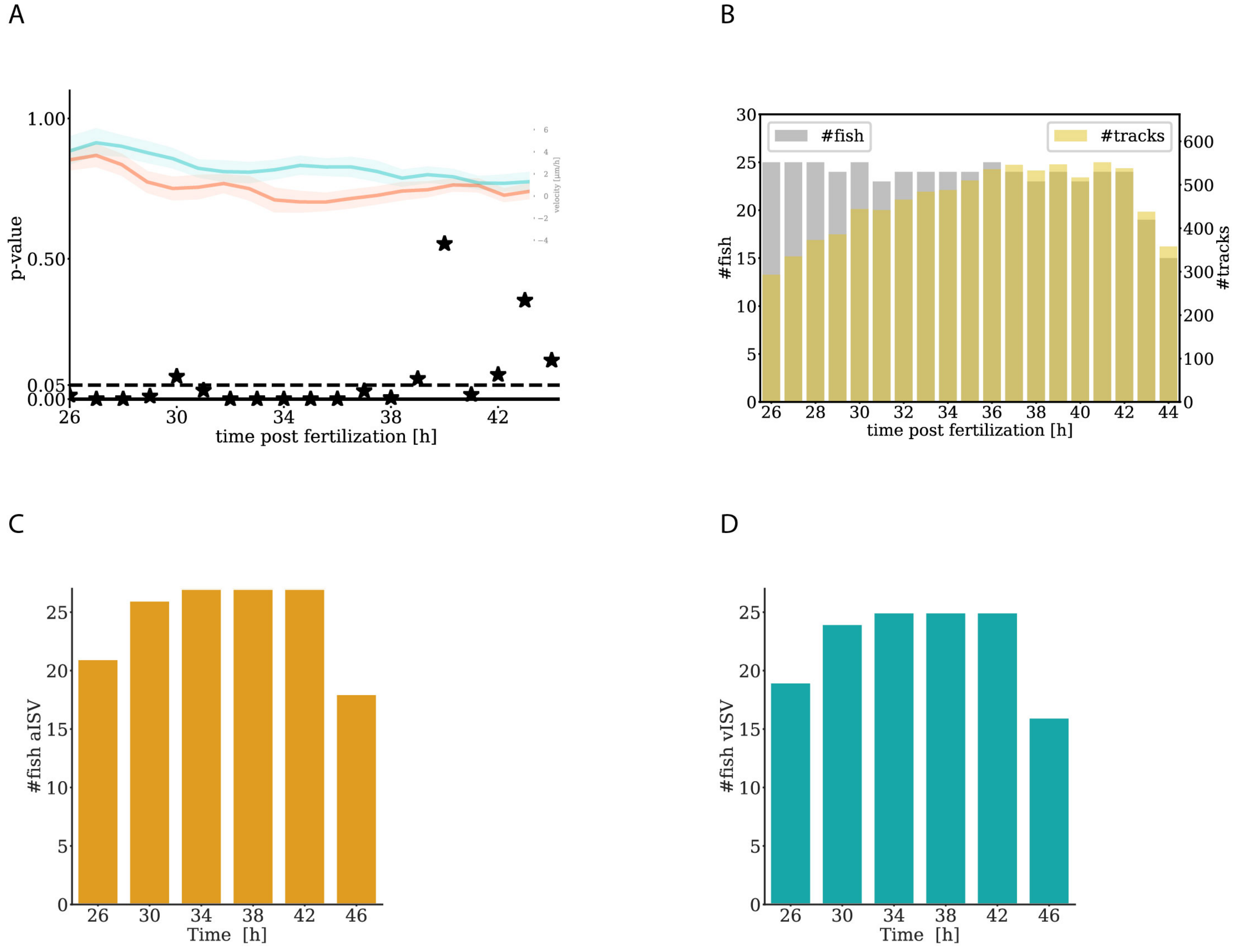

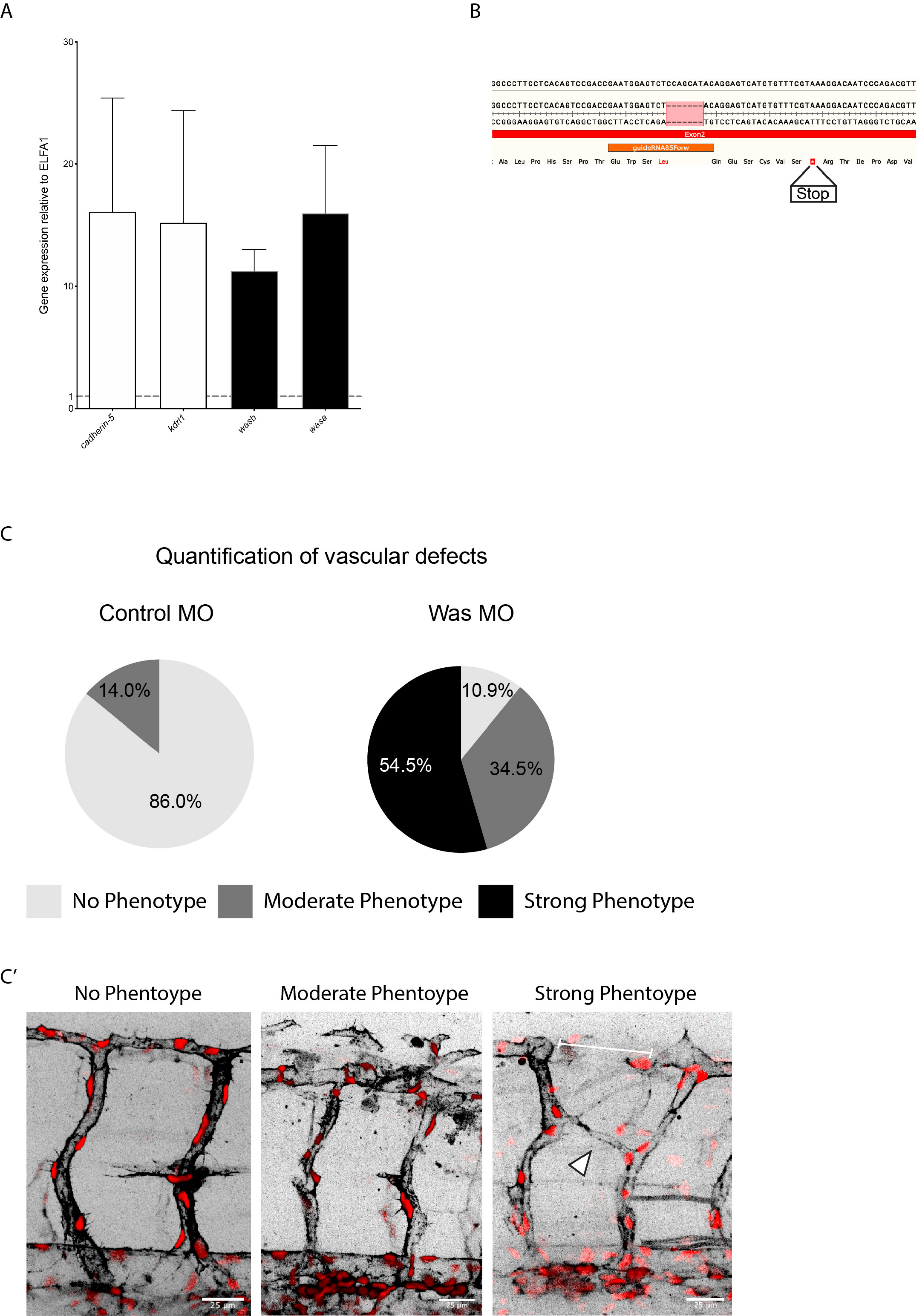

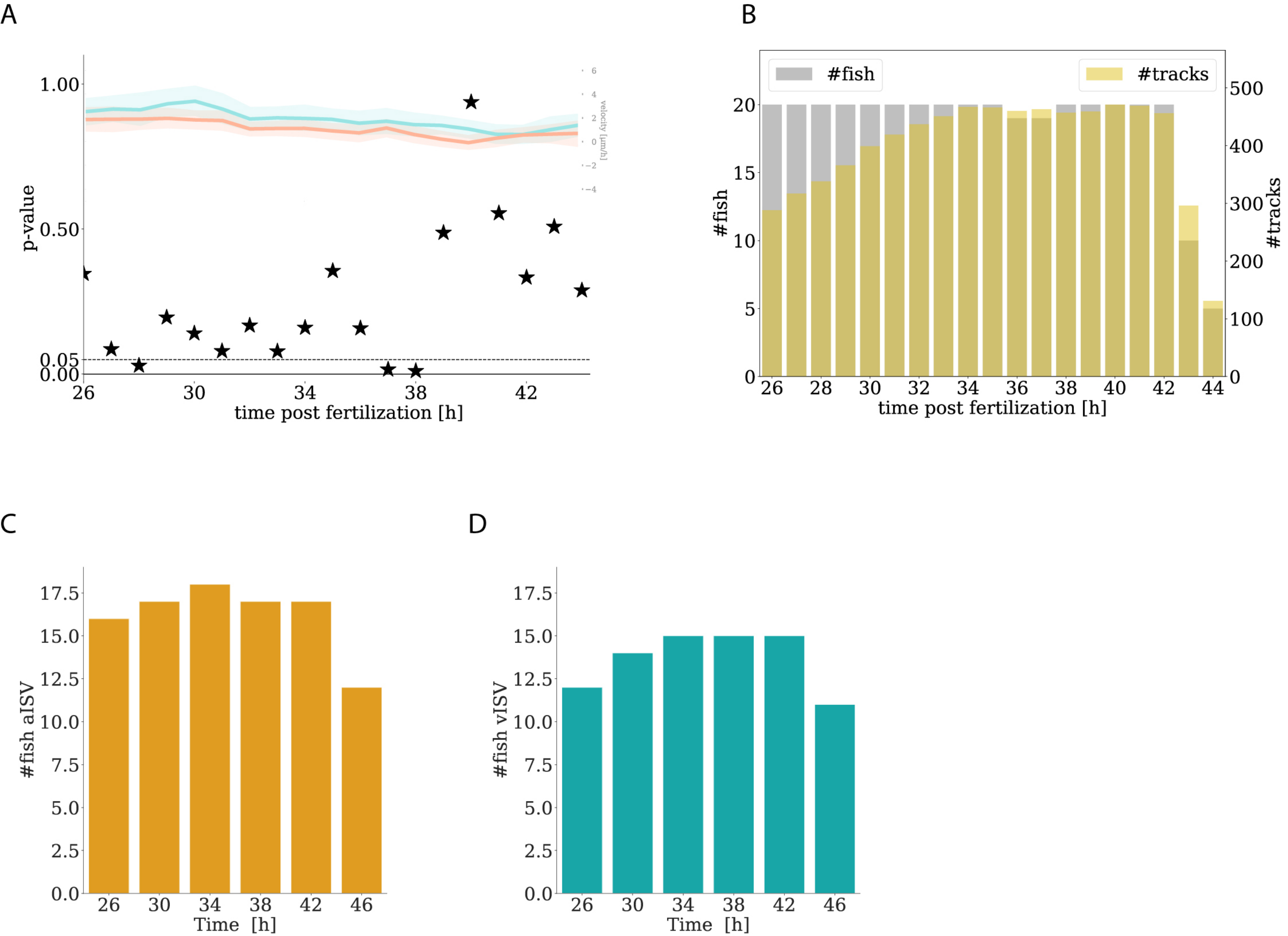

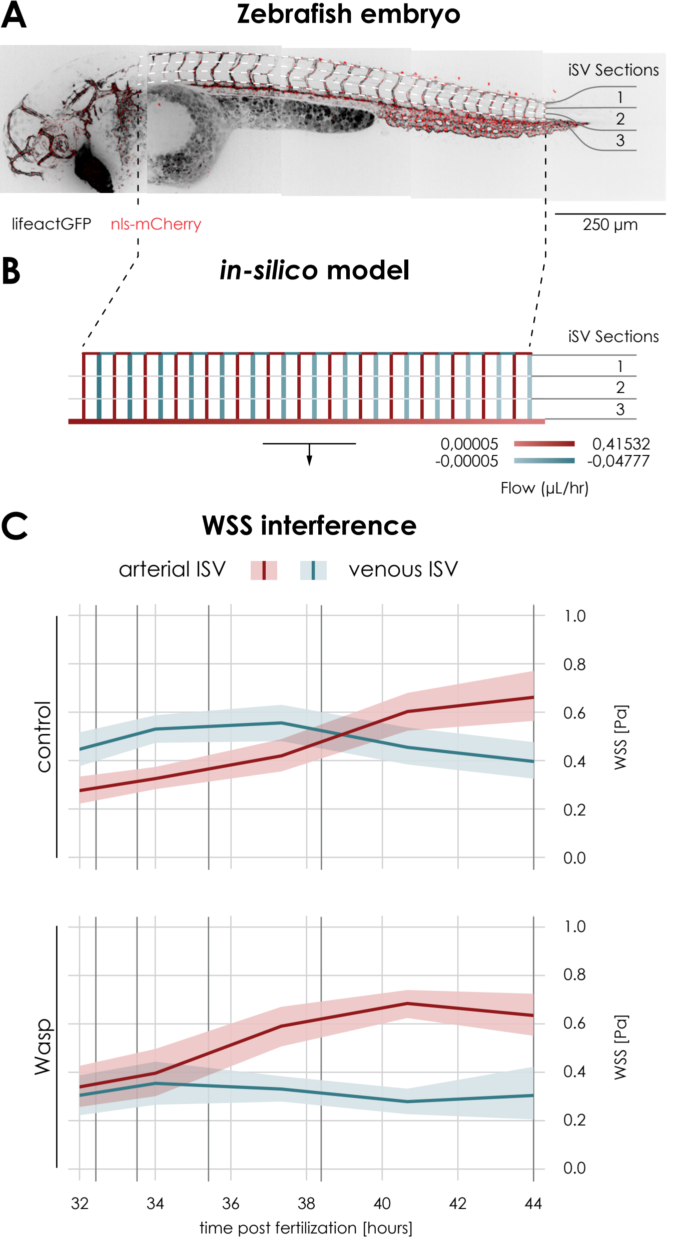

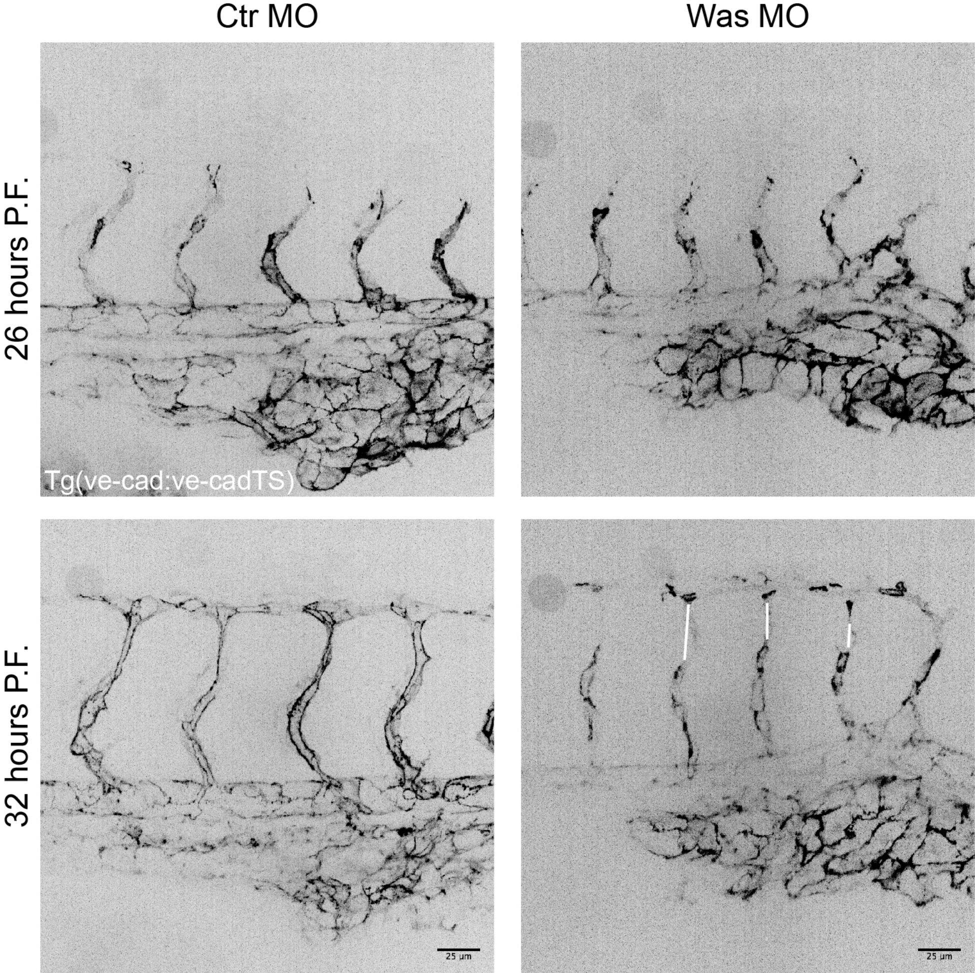

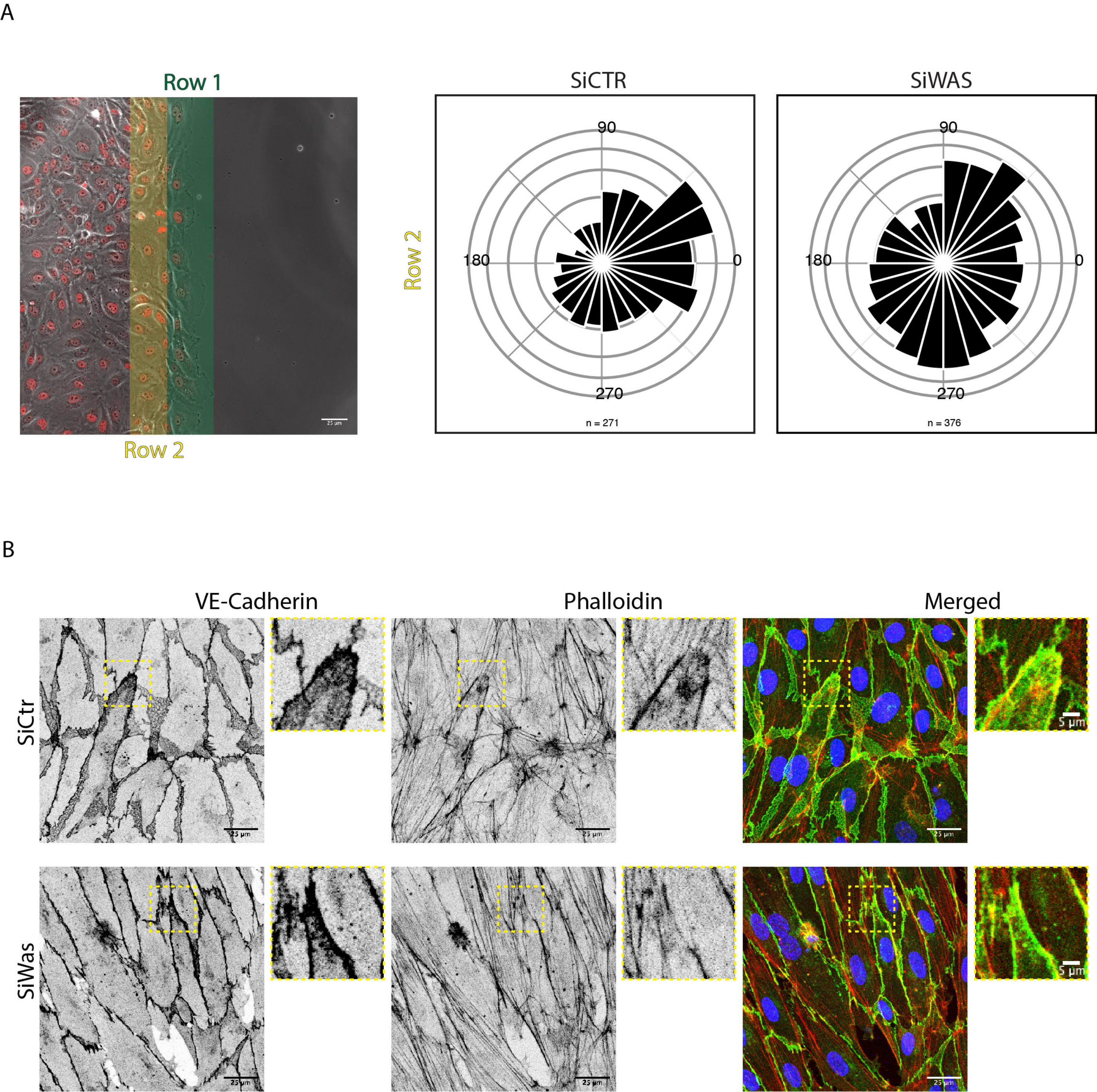
